# A machine learning framework for predicting and modulating condition-dependent protein phase separation

**DOI:** 10.64898/2025.12.28.696755

**Authors:** Jangwon Bae, Minjun Kang, Donghyuk Lee, Kuk-Jin Yoon, Yongwon Jung

**Affiliations:** Department of Chemistry, Korea Advanced Institute of Science and Technology, Daejeon, 34141, Republic of Koraea; The Robotics Program, Korea Advanced Institute of Science and Technology, Daejeon, 34141, Republic of Korea; Department of Mechanical Engineering, Korea Advanced Institute of Science and Technology, Daejeon, 34141, Republic of Korea

## Abstract

Protein phase separation is a fundamental process in organizing membraneless organelles and is implicated in a wide range of pathological conditions. Importantly, rather than being a static feature of specific proteins, phase separation is a condition-dependent phenomenon governed by environmental parameters, including protein concentration, temperature, and solvent composition. However, most existing machine learning models infer phase-separation propensity solely from amino-acid sequences, failing to capture these context-dependent behaviors. Here, we present LLPSense, a machine learning framework that integrates pre-trained protein language model embeddings with environmental parameters to achieve accurate, condition-aware predictions of protein phase separation. We demonstrate LLPSense’s predictive power and utility through three key experimental demonstrations. First, the model revealed that SGTA, previously unrecognized as a phase-separating protein, exhibits complex, temperature-dependent reentrant phase behavior. Second, LLPSense accurately predicted mutations in α-synuclein that either enhance or suppress phase separation, enabling systematic mapping of residues potentially relevant to Parkinson’s disease. Third, using model-guided mutagenesis, we inverted the phase behavior of UBQLN4, shifting it from high-temperature to low-temperature separation. Collectively, LLPSense provides a robust computational tool for interrogating the condition-dependent landscape of protein phase separation, enabling mechanistic studies of disease-associated phase separation and the rational design of programmable condensates.

## Introduction

Biomolecular phase separation of proteins and nucleic acids drives the formation of diverse membraneless organelles and contributes to the spatial organization of biochemical processes within cells^1^. Dysregulation of protein phase behavior has been increasingly implicated in the pathogenesis of various diseases, including neurodegenerative disorders^2–4^. Many proteins have been shown to undergo phase separation, thereby forming membraneless reaction hubs that regulate diverse cellular processes^5^. Despite substantial progress, experimentally identifying phase-separating proteins and characterizing their phase behavior remain labor-intensive and low-throughput. To overcome these limitations, numerous machine learning (ML) approaches have been developed. In particular, several ML predictors^6–9^ have been trained to estimate phase separation propensity directly from protein sequences, achieving superior performance compared with earlier *in silico* screening methods^10^. Among them, the PSPire^11^ exhibits remarkable predictive accuracy, even for proteins lacking intrinsically disordered regions (IDRs). Another ML model, PSPHunter^12^, identified key residues that drive phase separation and demonstrated that truncation of 6 residues in GATA3 was sufficient to inhibit its phase separation. More recently, catGRANULE 2.0^13^ has shown improved predictive performance relative to these predecessors and was the first to attempt to predict mutants with increased phase-separation propensity.

Previous sequence-based ML prediction models have relied on static binary classification (i.e., whether a protein is capable of phase separation). However, natural liquid-liquid phase separation (LLPS) of proteins is a dynamic, stimulus-responsive phenomenon governed by complex physicochemical conditions (e.g., pH, temperature)^14, 15^. Because conventional methods rely exclusively on sequence feature similarity, they inherently treat phase separation as a fixed property, failing to account for the fact that the same protein’s behavior varies with conditions. This reliance limits their ability to precisely estimate relative LLPS propensity, which is vital for characterizing the divergent effects of mutations that can either suppress or enhance phase behavior. Therefore, capturing the full spectrum of such modulation requires bidirectional scales that enable precise comparison of phase separation probabilities, supported by a comprehensive dynamic range spanning the global condition space. Droppler^16^ represents a pioneering effort to address this limitation by employing a neural network to estimate LLPS probability under various conditions. Despite this advancement, its predictive performance remains limited, thereby restricting broader applicability. The authors of Droppler acknowledged that modeling condition-dependent LLPS is substantially more challenging than sequence-only classification. Consequently, developing a robust predictor that integrates both intrinsic protein sequences and extrinsic conditional variables is essential for the scalable computational analysis of LLPS behavior across large protein datasets^15^. Such a condition-aware framework would provide a powerful platform for investigating LLPS biology and for enabling the *in silico* design of programmable biomolecular condensates.

In this study, we present LLPSense, a condition-aware ML framework for predicting protein phase behavior across diverse physicochemical environments. LLPSense accurately estimates the phase-separation probability of a given protein sequence under a wide range of parameters, including protein concentration, temperature, pH, crowding agents, salt concentration, and glycerol content. We thoroughly validated the predictive power and experimental utility of LLPSense through multiple analyses. Initially, we challenged the model to screen for overlooked phase-separating proteins within datasets previously classified as negative. LLPSense flagged 16.4% of these sequences as potential phase-separating candidates, and this predicted subset included proteins already reported in the literature to undergo phase separation. Notably, this screen identified SGTA as a previously unrecognized LLPS protein, whose complex reentrant, temperature-dependent phase behavior was accurately predicted and subsequently confirmed experimentally. LLPSense also pinpointed specific residues in Parkinson’s disease-associated α-synuclein, predicting substitutions that either enhance or suppress phase separation. Comparative analysis with a sequence-only predictor confirmed that the condition-aware framework facilitates mutational analysis at single-residue resolution and bidirectional prediction. We additionally demonstrated that the conditional dependence of LLPS can be modulated by altering intrinsic sequence properties. Specifically, mutational engineering of the lower critical solution temperature (LCST)-type protein UBQLN4 shifted its phase behavior toward an upper critical solution temperature (UCST)-like regime, highlighting that LLPSense can guide the rational design of sequence modifications for tuning condition-specific phase behavior.

## Results

### Protein language model embeddings facilitate accurate prediction of phase separation

Motivated by the recent success of applying language-based encoding methods, such as Word2Vec, to LLPS prediction^7, 12^, we hypothesized that leveraging advanced pretrained protein language models (pLMs) could provide a more accurate and efficient framework. Among available pLMs, we selected ProtT5^17^ to extract high-quality sequence embeddings for both training and inference, as prior work (Seq2Phase)^18^ has demonstrated that pLM-derived embeddings enable accurate prediction of proteins recruited to biomolecular condensates. Before integrating environmental variables, we validate this pLM-based strategy using the conventional sequence-based, binary LLPS prediction task. We selected PSPire as a comparative reference because, in addition to its high predictive performance near the completion of our model, it demonstrated improved identification of phase-separating proteins lacking intrinsically disordered regions (IDRs). This distinction is crucial for assessing whether pLMs can provide generalized protein representations that capture phase separation properties across the full structural spectrum. Accordingly, we trained and evaluated our models using the PSPire dataset, which categorizes human proteins into three groups: intrinsically disordered phase-separating proteins (ID-PSPs), structured phase-separating proteins (noID-PSPs), and non-phase-separating proteins (non-PSPs). For binary classification, ID-PSPs and noID-PSPs were treated as positive samples, while non-PSPs were considered negatives.

We developed a gradient-boosted decision tree classifier based on XGBoost^19^. A previous study demonstrated that tree-based classifiers, such as XGBoost and Random Forest, outperform neural-network architectures and other ML approaches for LLPS prediction^12^. We first obtained pLM-based embeddings using ProtT5, which encodes an amino acid sequence of length N into an N × 1,024 embedding matrix. Mean pooling was then applied across the sequence dimension to obtain a single 1,024-dimensional vector representation. These averaged embeddings were subsequently used as input features for model training and evaluation, yielding a model we termed LLPSeq (Fig. 1a). This model demonstrated remarkable predictive performance. In both the ID and noID test sets, the area under the receiver operating characteristic curve (AUROC) was comparable to values reported for PSPire, while the area under the precision-recall curve (AUPRC) surpassed it (Fig. 1b–e). To benchmark protein language embeddings against conventional feature-based models with a similar architecture, we constructed a baseline model trained on 55 knowledge-based engineered features (e.g., molecular weight, net charge, hydrophilicity, and low-complexity region content) widely used in previous studies^6, 7^. The overall performance of this baseline model was inferior to that of PSPire and LLPSeq, especially in the IDR-lacking (noID) test set (Fig. 1d, e). These findings indicate that, even without handcrafted feature engineering, embeddings derived directly from pretrained protein language models provide a more effective and generalizable representation for LLPS prediction.

**Figure 1.**
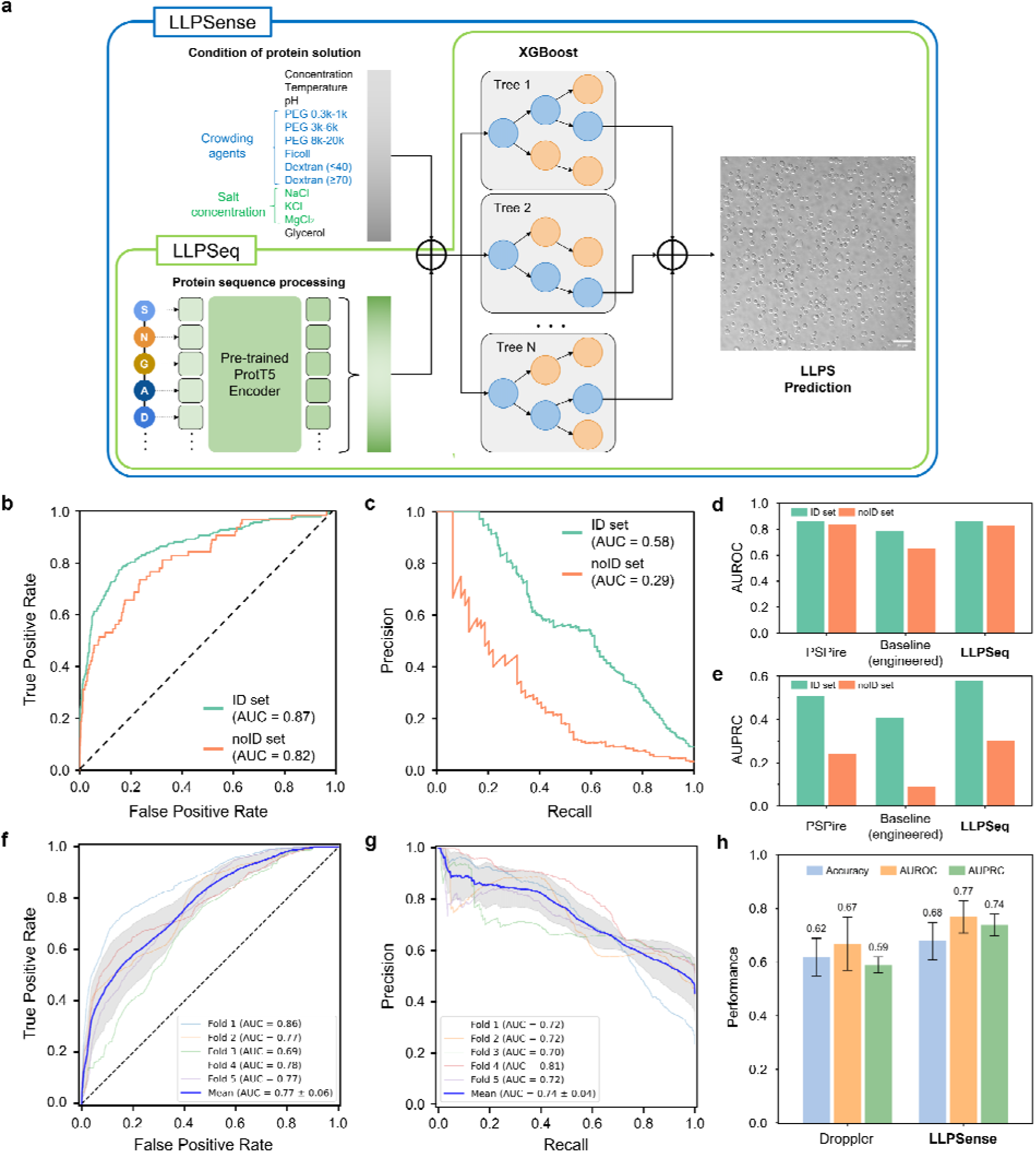
Architectural framework and performance benchmarking of LLPSeq and LLPSense. **(a)** Schematic illustration of the LLPSeq and LLPSense frameworks. **(b, c)** Predictive performance of the pLM (protein language model)-based LLPSeq model, shown as (b) receiver operating characteristic (ROC) curves and (c) precision-recall (PR) curves. Summary of **(d)** AUROC and **(e)** AUPRC scores of LLPSeq against PSPire and a baseline model (trained on 55 engineered features). **(f, g)** Evaluation of LLPSense via 5-fold cluster-based cross-validation, displaying (f) ROC and (g) PR curves. Shaded regions represent ± 1 s.d. from the mean. **(h)** Performance comparison on the sequence-condition integrated task against the Droppler model, which was retrained on the same expanded dataset. Error bars represent ± 1 s.d. from 5-fold cross-validation.

### LLPSense accurately predicts condition-dependent protein phase behavior

Building on the promising performance of the pLM-based LLPSeq model, we extended this framework to develop a comprehensive predictor that integrates both intrinsic sequence information and extrinsic environmental conditions. A significant bottleneck in developing condition-aware models has been the limited availability of large, standardized datasets suitable for machine learning. Although LLPSDB v2^20^ provides a broad collection of *in vitro* LLPS observations, substantial curation is required to render it amenable to model training. Therefore, we assembled a rigorously curated, ML-ready dataset by filtering LLPSDB v2 specifically for single-protein entries.

For each entry, we extracted and standardized a comprehensive set of 13 condition variables, including protein concentration, temperature, pH, six representative categories of crowding agents, concentrations of three representative salts (NaCl, KCl, and MgCl_), and glycerol percentage. These 13 variables were sufficient to capture the majority of experimentally annotated entries. In contrast, complex cosolutes such as heparin or RNA were excluded due to data sparsity and mechanistic complexity. Collectively, these selected parameters capture key physicochemical modulators of LLPS, encompassing solvent polarity, ionic strength, and macromolecular crowding. Furthermore, to accommodate the continuous nature of experimental phase diagrams, we implemented a systematic interval-partitioning strategy. This method subdivides reported ranges into uniformly spaced samples, thereby augmenting the effective number of training instances while preserving experimental validity (see Methods for details). All curated datasets and preprocessing scripts used in this study are publicly available to facilitate future machine learning research on condition-dependent protein phase separation (see Data Availability).

Next, sequence embeddings derived from ProtT5 were concatenated with the normalized condition variables to construct an integrated feature matrix. This combined representation served as the input for an XGBoost-based architecture, and we designated this predictive framework as LLPSense (Fig. 1a). This multimodal integration enables the model to simultaneously capture intrinsic sequence determinants of LLPS and extrinsic physicochemical factors that modulate condensate formation. We employed a 5-fold cluster-based cross-validation (CV) strategy to assess the generalizability of LLPSense and identify optimal hyperparameters. To prevent overestimation of performance due to sequence homology between training and test sets, proteins were grouped into 234 distinct clusters using MMseqs2^21^, ensuring that sequences in different clusters shared <20% identity at 90% coverage. All proteins within the same cluster were assigned to the same CV fold, thereby maintaining strict separation between training and validation sets. Under this cluster-based 5-fold CV, LLPSense demonstrated robust predictive performance, achieving an average AUROC of 0.77 and an AUPRC of 0.74 (Fig. 1f–h). This performance significantly surpasses the AUROC value reported for Droppler (0.64). Furthermore, even when Droppler was retrained on our expanded dataset, LLPSense continued to exhibit superior predictive accuracy (Fig. 1h, Supplementary Fig. 1). Altogether, these results demonstrate that our integrated modeling strategy provides an effective and reliable framework for predicting condition-dependent phase separation. Given that modeling conditional dependencies is significantly more challenging than sequence-only prediction, the achieved performance marks a pivotal milestone in the field. We next investigated whether this *in silico* predictive accuracy translates into practical utility in real experimental settings.

### Identification of potential phase-separating proteins in the previously classified negative dataset

We screened the human proteome dataset previously classified as non-PSP (used in the PSPire study) using LLPSense. Remarkably, a substantial fraction of proteins (16.4%, 1,681/10,252) were predicted to undergo phase separation within a temperature range of 10–50 °C under physiological conditions (100 µM protein concentration, pH 7.3, and 160 mM NaCl; Fig. 2a). Among these proteins, SS18^22^, FOXP2^23^, and Atrophin-1^24^ (Fig. 2b) have been reported to exhibit phase-separating behavior, despite their inclusion in the non-PSP set. This discrepancy underscores a critical limitation in the existing negative datasets. Such issues arise primarily because a definitive list of proteins that are strictly incapable of LLPS has not been established. As a result, previous studies have resorted to heuristic negative sets derived from indirect criteria, such as subtracting known LLPS-positive proteins from the proteome. However, the boundary between “LLPS-competent” and “LLPS-incompetent” proteins is inherently fluid, varying across environmental conditions and experimental methods. This ambiguity complicates model training and further emphasizes the need for condition-aware strategies.

**Figure 2.**
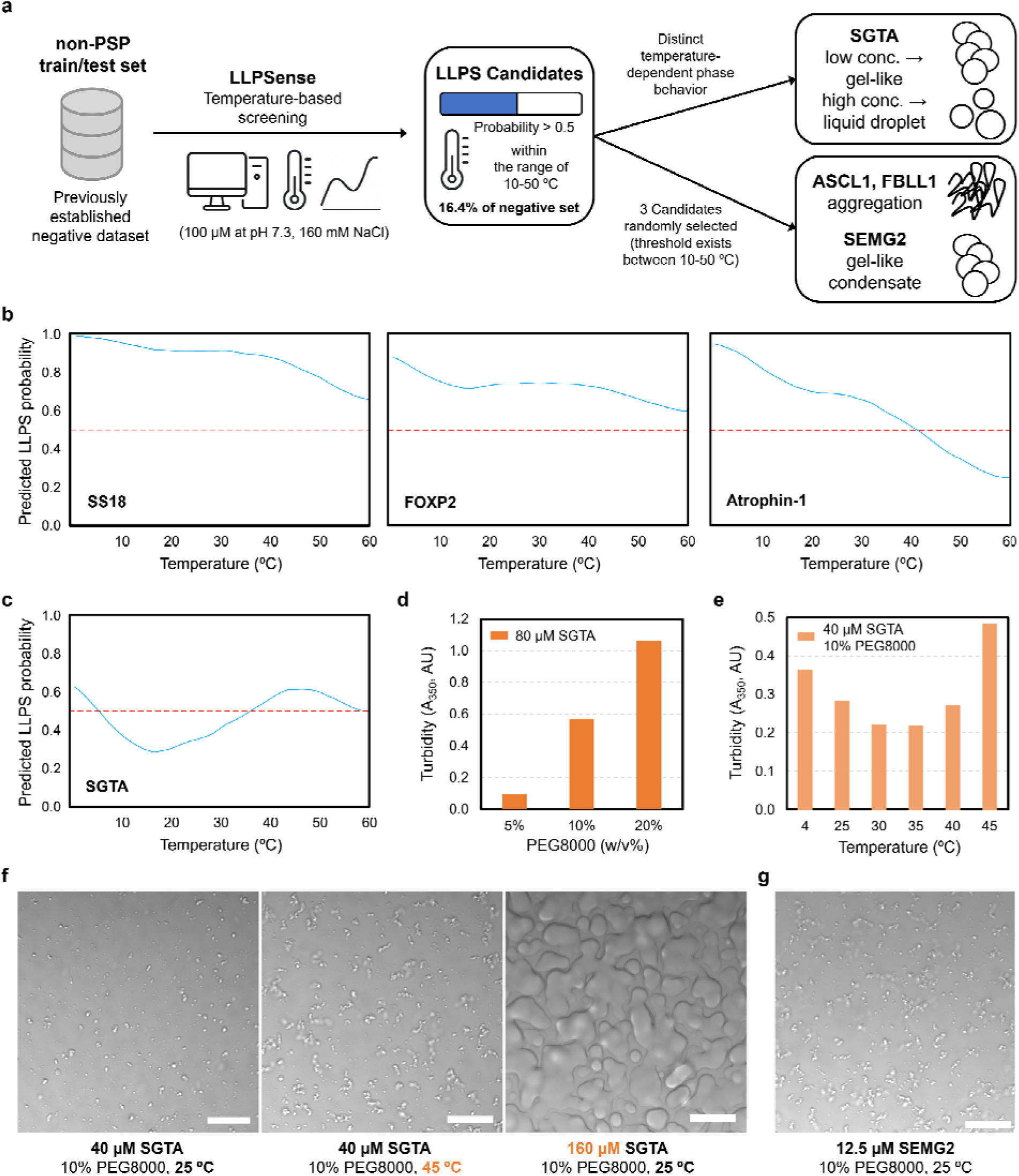
Experimental validation of LLPS candidates identified by LLPSense from a previously defined negative dataset. **(a)** Schematic overview of the screening and validation workflow. LLPSense re-evaluated a previously established non-PSP dataset, identifying 16.4% of entries as potential phase separation candidates (probability > 0.5 within 10–50 °C). **(b, c)** Predicted temperature profiles of LLPS candidates. The curve was smoothed using a centered moving average (window size = 15). (b) Literature-validated phase separating proteins previously misclassified in the non-PSP set. (c) The SGTA protein profile exhibits distinctive reentrant phase behavior, characterized by mixed UCST/LCST regimes. **(d)** Turbidity measurements (A_350_) for SGTA under crowding conditions. **(e)** Temperature-dependent turbidity validating the predicted reentrant behavior of SGTA. **(f)** Representative microscopy images showing the condition-dependent morphology of SGTA condensates. **(g)** SEMG2 condensates exhibiting interconnected gel-like clusters. (f, g) Scale bars, 20 μm.

Among the candidates newly identified by LLPSense, SGTA stood out for its predicted temperature-dependent reentrant phase behavior, which displays characteristics of both the upper critical solution temperature (UCST) and lower critical solution temperature (LCST) regimes (Fig. 2c). To experimentally validate this prediction, SGTA was recombinantly expressed, and its phase behavior was assessed using solution turbidity measurements, a commonly used quantitative indicator of phase-separation propensity. We observed a PEG concentration-dependent increase in SGTA turbidity at 350 nm (A_350_), confirming its intrinsic ability to phase separate (Fig. 2d). Consistent with the predicted reentrant profile, turbidity increased modestly at 30–35 °C but showed a much greater increase at both low (4 °C) and elevated (≥ 40 °C) temperatures (Fig. 2e). Microscopic images of SGTA condensates corroborated these findings, showing that the total volume of condensates was larger at elevated temperatures (Fig. 2f). Collectively, these results demonstrate that LLPSense not only identifies previously unrecognized phase-separating candidates but also captures their intricate, condition-dependent phase behavior.

To further assess the predictive applicability of LLPSense, we randomly selected three proteins (ASCL1, FBLL1, SEMG2) from candidates predicted to exhibit a phase separation threshold within the 10–50 °C range (Supplementary Fig. 2) and evaluated their behavior *in vitro*. ASCL1 and FBLL1 aggregated during dialysis against PBS (pH 7.3, 160 mM NaCl), resulting in final concentrations below 10 μM. FBLL1 is known to localize within the nucleolus^25^, suggesting that it may participate in the phase behavior of this membraneless organelle. SEMG2 is a semenogelin protein responsible for the gelation of seminal fluid^26^. Upon addition of PEG8000, SEMG2 displayed marked turbidity (Supplementary Fig. 3), consistent with macromolecular condensation.

The aggregation of ASCL1 and FBLL1 may reflect their high intrinsic LLPS propensity, as phase-separating proteins often exhibit marginal solubility and a greater tendency to aggregate^27, 28^. Furthermore, we observed a concentration-dependent transition in the material state: SGTA formed large liquid-like droplets at 160 μM (with 10% PEG8000), whereas small, gel-like condensates formed at lower concentrations (Fig. 2f). SEMG2 also formed interconnected gel-like condensates rather than canonical liquid droplets (Fig. 2g). Despite extensive research, the criteria for distinguishing liquid-like condensates from percolated gels and irreversible aggregates remain poorly defined^29^. Accordingly, while LLPSense effectively predicts condensation propensity, it does not yet discriminate among distinct material states. With the expanding understanding of biomolecular phase behavior, LLPS, gelation, aggregation, and fibrillization are viewed not as discrete end states but as interconnected regimes within a shared energy landscape^30–32^. Our preliminary tests with ASCL1, FBLL1, SEMG2, and SGTA illustrate this continuum, highlighting both the broad predictive reach of LLPSense and the necessity for future data refinements that account for the full liquid–gel–solid spectrum.

### *In silico* mapping of the α-synuclein mutational landscape at single-residue resolution

Many diseases are associated with mutations that induce aberrant enhancement^33–36^ or suppression^37, 38^ of phase separation. However, relying solely on sequence information poses a significant risk of overfitting to limited positive datasets, which often lack environmental context. Furthermore, such approaches may struggle to capture the relative phase-separating capacity of different sequences, as all proteins validated to undergo LLPS are typically treated as uniform positive instances. Our condition-aware framework addresses these limitations by providing the granularity needed to distinguish sequence behavior under varying environments. To illustrate this capability, we examined α-synuclein, whose LLPS behavior is increasingly recognized as a critical intermediate state preceding pathological fibrillization^39^. By systematically substituting each of the 140 residues of α-synuclein and computing the resulting changes in LLPS probability, we predicted specific mutations that modulate its phase-separation landscape.

LLPSense enabled rapid probability screening across the entire mutational landscape, comprising all possible single-point substitutions (140 residues × 19 variants; Fig. 3a). To experimentally validate the predicted effects, we selected two mutations projected to exert the strongest influence on phase separation (Fig. 3b). Specifically, the substitution of glutamate with methionine at position 28 (E28M) within the amphipathic N-terminal domain yielded the highest predicted increase in LLPS probability; we termed this the ‘E1 variant’. Conversely, the substitution of valine with aspartate at position 77 (V77D) within the hydrophobic non-amyloid component (NAC) domain resulted in the most significant reduction, designated the ‘S1 variant’. By comparison, mutations within the acidic C-terminal tail elicited relatively minor fluctuations in predicted phase behavior.

**Figure 3.**
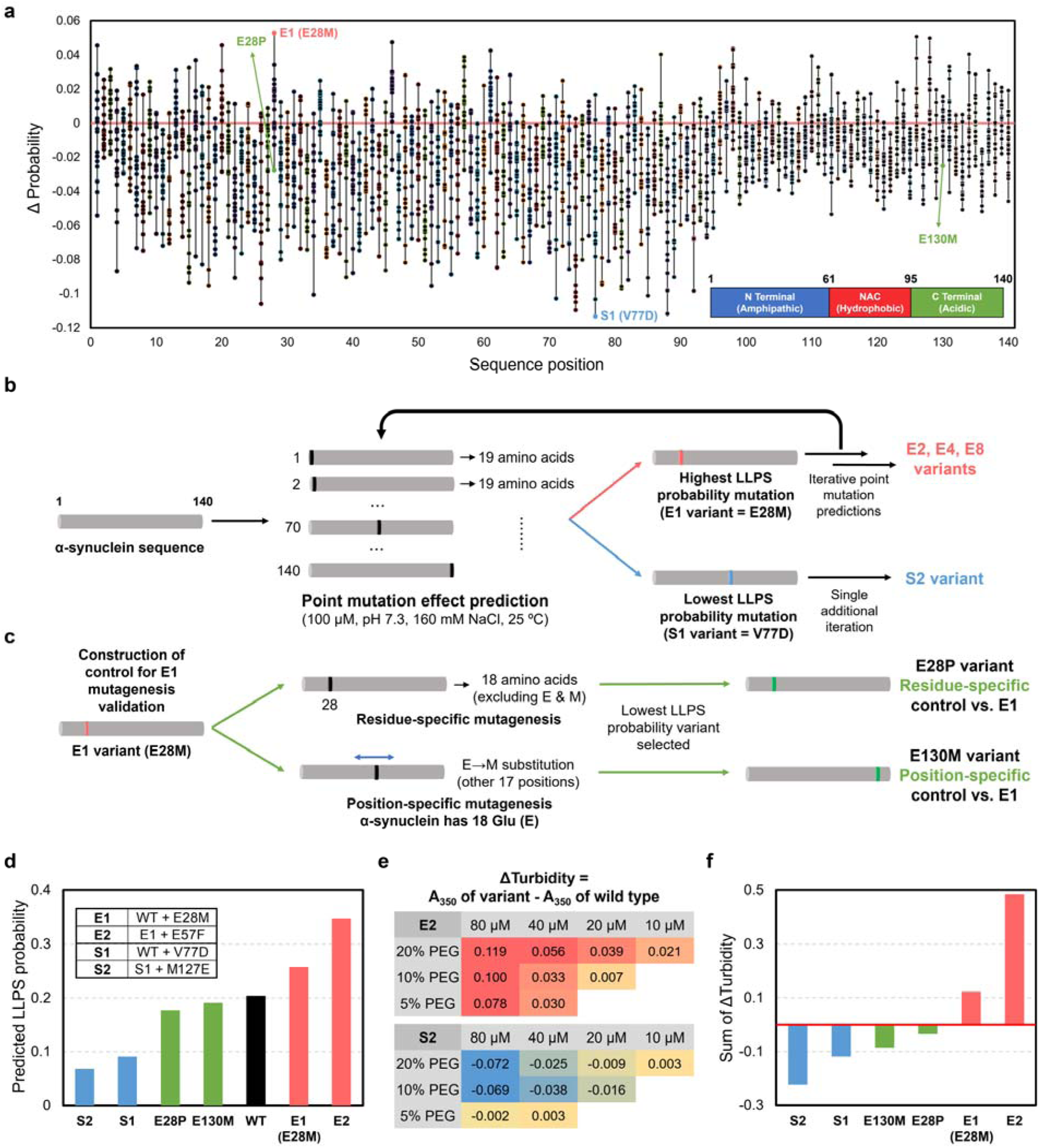
Residue-level LLPS propensity mapping of α-synuclein variants using LLPSense. **(a)** Predicted mutational landscape showing the shift in probability (ΔProbability) for all point mutants relative to the WT (red line). Key variants (E1, S1) and controls (E28P, E130M) are highlighted. The domain architecture of α-synuclein is annotated at the bottom. **(b)** Schematic workflow of the *in silico* saturation mutagenesis. E1 (E28M) and S1 (V77D) were identified as the strongest modulators, serving as scaffolds for recursive multi-mutant design (E2–E8, S2). **(c)** Schematic of control variant construction to verify the specificity of LLPSense. Controls were designed to decouple position and residue effects: E28P (same position, different residue) and E130M (different position, same residue). **(d)** Predicted probabilities for selected variants. The inset details the specific mutations. **(e)** Experimental ΔTurbidity heatmaps showing condition-dependent phase behavior relative to WT. Warm colors indicate enhancement; cool colors indicate suppression. **(f)** Summed ΔTurbidity across all conditions. The experimental ranking mirrors the predictions in (d), verifying the distinct separation of enhanced and suppressed variants relative to the WT baseline (red line).

Using the E1 and S1 variants as scaffolds, we performed iterative rounds of prediction to identify additional mutations that confer additive effects while retaining previously mutated positions. Through this recursive procedure, we constructed a series of variants with multiple mutations, designating the phase-separation enhanced set as E2, E4, and E8, and the suppressed set as S2 (Supplementary Table 1). For these variants, the numerical suffix indicates the cumulative number of incorporated mutations. In addition, to verify that LLPSense accurately predicted E1 based on the specific position (E28) and residue (E-to-M), two control mutants (E28P and E130M) were selected for their lowest predicted LLPS probabilities among possible combinations (Fig. 3c). E28P represents a different substitution at the same position, while E130M represents the identical E-to-M substitution at a different position. Both control mutants exhibit predicted probability significantly lower than that of the E1 variant and slightly below the WT baseline (Fig. 3d).

We expressed and purified all α-synuclein variants under identical conditions and assessed their phase separation behavior by measuring turbidity across nine combinations of PEG8000 and protein concentrations. Heatmap visualization of ΔTurbidity, calculated by subtracting WT turbidity from that of each variant’s turbidity, revealed a stark contrast between the enhanced and suppressed variant sets (Fig. 3e, Supplementary Fig. 4). Summing the ΔTurbidity values across all nine conditions yielded the following order: E2 > E1 > WT (baseline) > E28P > E130M > S1 > S2 (Fig. 3f). This ranking was in excellent agreement with LLPSense predictions (Fig. 3d), with the only minor deviation occurring between E130M and E28P, where predicted differences were marginal. Consistent with these results, microscopic analysis of the α-synuclein variants confirmed that those with higher predicted propensities exhibited a clear increase in condensate formation (Supplementary Fig. 5). During PBS dialysis, the hyper-enhanced variants E4 and E8 exhibited pronounced aggregation and structural instability (Supplementary Fig. 6a, b). This observation corroborates our earlier discussion that condensation-prone sequences often exhibit a concomitant increase in aggregation propensity. Nonetheless, both variants still showed turbidity levels higher than WT (Supplementary Fig. 6c). Taken together, these results demonstrate that LLPSense accurately identifies mutations that modulate phase separation and can reliably pinpoint sequence alterations that shift α-synuclein toward LLPS-prone states.

### Condition-aware mutational screening affords a bidirectional scale and granular resolution of LLPS propensity

To demonstrate the enhanced accuracy and granularity of our condition-aware framework, we also compared the mutational landscapes of α-synuclein predicted by LLPSense and the sequence-based predictor LLPSeq. Despite the absence of α-synuclein from its training dataset, LLPSeq predicted a high probability of 0.78 for WT, a result likely driven by overfitting to the homologous β-synuclein sequence. Consequently, LLPSeq classified most single-point mutations as suppressive (90.2%; 2,039/2,260). In contrast, even with the target protein included in its training, LLPSense identified a substantial fraction of mutations as enhancing (24.0%; 543/2,260) (Fig. 4a, b), demonstrating a broad bidirectional prediction range. LLPSeq also predicted a monotonic decrease in phase separation potential for enhanced mutants (E1–E8), showing an increasingly pronounced discrepancy from the intended cumulative modulation (Supplementary Fig. 7). In addition, LLPSense can circumvent this saturation issue through its environmental sensitivity. By modulating parameters such as protein concentration or crowding agent levels, the model can shift the baseline into a neutral regime, near a predicted probability of 0.5, even if a sequence exhibits high intrinsic probability. This capability ensures a sufficient dynamic range to resolve subtle mutational effects that are otherwise masked in static, sequence-only models.

**Figure 4.**
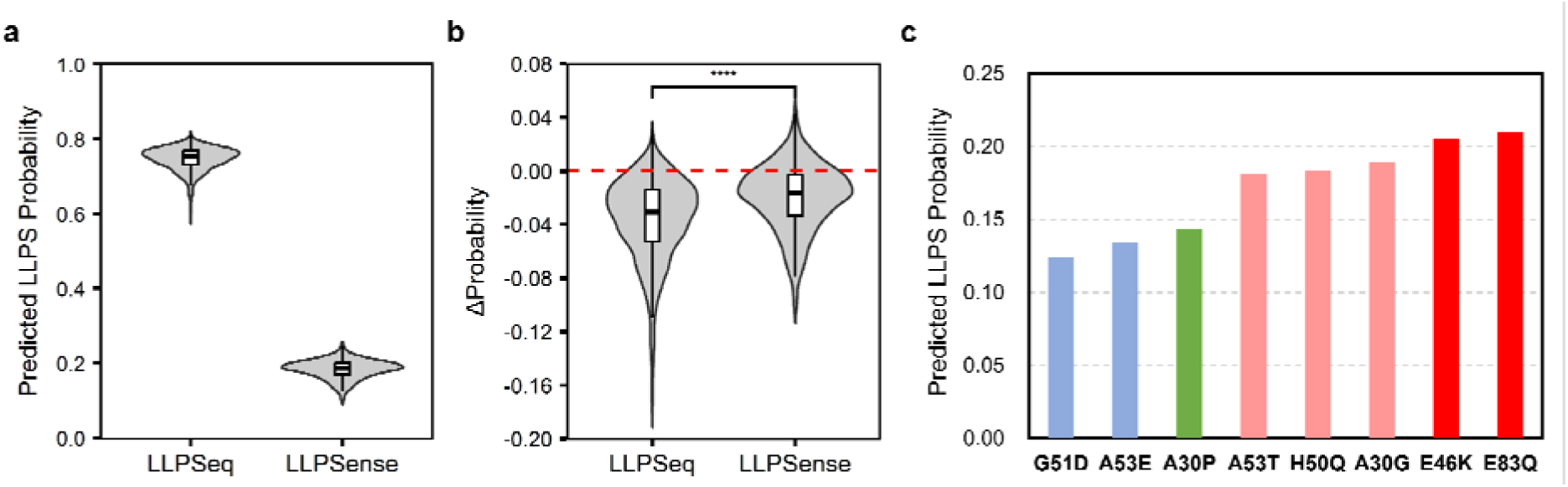
Condition-aware screening resolves the bidirectional and granular mutational impacts on α-synuclein LLPS. **(a)** Distribution of predicted LLPS probabilities for the α-synuclein single-point mutation library. **(b)** Comparison of the mutational landscape between LLPSeq and LLPSense. The violin plots show the distribution of ΔProbability (the change in probability relative to the wild-type). Statistical significance was determined by a two-sided Wilcoxon rank-sum test (****PL<L0.0001). Boxplots represent the interquartile range (IQR) from Q1 to Q3, with the median as the middle line. Whiskers extend up to 1.5 times the IQR. Outliers are not shown. **(c)** The variants are color-coded based on experimental LLPS propensity measurements reported in the literature: red (significantly enhanced), pink (mildly enhanced), green (comparable), and blue (suppressed) compared to WT. The predicted probabilities show a high qualitative correlation with the reported experimental rankings: (1) E46K ≈ E83Q > A30G ≈ A53T > A30P ≥ WT > A53E > G51D and (2) E46K > H50Q ≈ A53T > WT ≈ A30P > A53E (3) the identification of E46K, A53T, and H50Q as LLPS-enhancing mutations.

We additionally benchmarked LLPSense on α-synuclein variants previously reported as pathogenic and characterized for LLPS propensity using turbidity and droplet-size measurements^39–42^. Based on prior experimental observations, these mutations were categorized into four distinct groups: significantly enhanced, mildly enhanced, comparable to WT, and suppressed. LLPSense demonstrated high qualitative consistency with these experimental groupings by accurately reflecting the relative rankings and propensity shifts across all tested substitutions (Fig. 4c). The absolute probability for the WT sequence (0.204) may be subject to moderate overestimation due to its inclusion in the training set, though this is expected to be mitigated as the dataset’s diversity and size increase. Nevertheless, the relative comparisons and predicted trends across various mutants remain highly valid, demonstrating that LLPSense accurately reflects the impact of each substitution at single-residue resolution. In contrast, while an existing bidirectional mutation model, catGRANULE 2.0, captured general tendencies, it incorrectly predicted the relative order of certain variants (Supplementary Fig. 8). It also predicted trends for our discovered S1 and E1 variants that were opposite to the experimental observations, which illustrates the difficulty of accurately predicting subtle mutational impacts (Supplementary Table 2).

### LLPSense-guided mutations reprogram UBQLN4 from LCST to UCST phase behavior

Most phase-separating proteins exhibit upper critical solution temperature (UCST) behavior, where phase separation is favored at lower temperatures due to reduced entropic costs. In contrast, certain families, such as elastin-like polypeptides (ELPs)^43^ and the UBQLN family^44^, exhibit LCST behavior, undergoing phase separation as temperature increases. Our curated dataset included ELPs and UBQLN2, but excluded UBQLN1 and UBQLN4. Notably, while UBQLN1 shares high sequence homology with UBQLN2 (76.21% identity, 82.79% similarity), UBQLN4 is more distantly related (58.72% identity, 70.19% similarity). When the Droppler model was retrained on our dataset, it correctly classified the highly homologous UBQLN1 as an LCST-type protein but failed to identify the more divergent UBQLN4 (Supplementary Fig. 9a). By contrast, LLPSense accurately predicted the LCST behavior of both UBQLN1 and UBQLN4 (Fig. 5a), demonstrating improved generalizability and sensitivity to the sequence determinants underlying temperature-dependent phase behavior.

**Figure 5.**
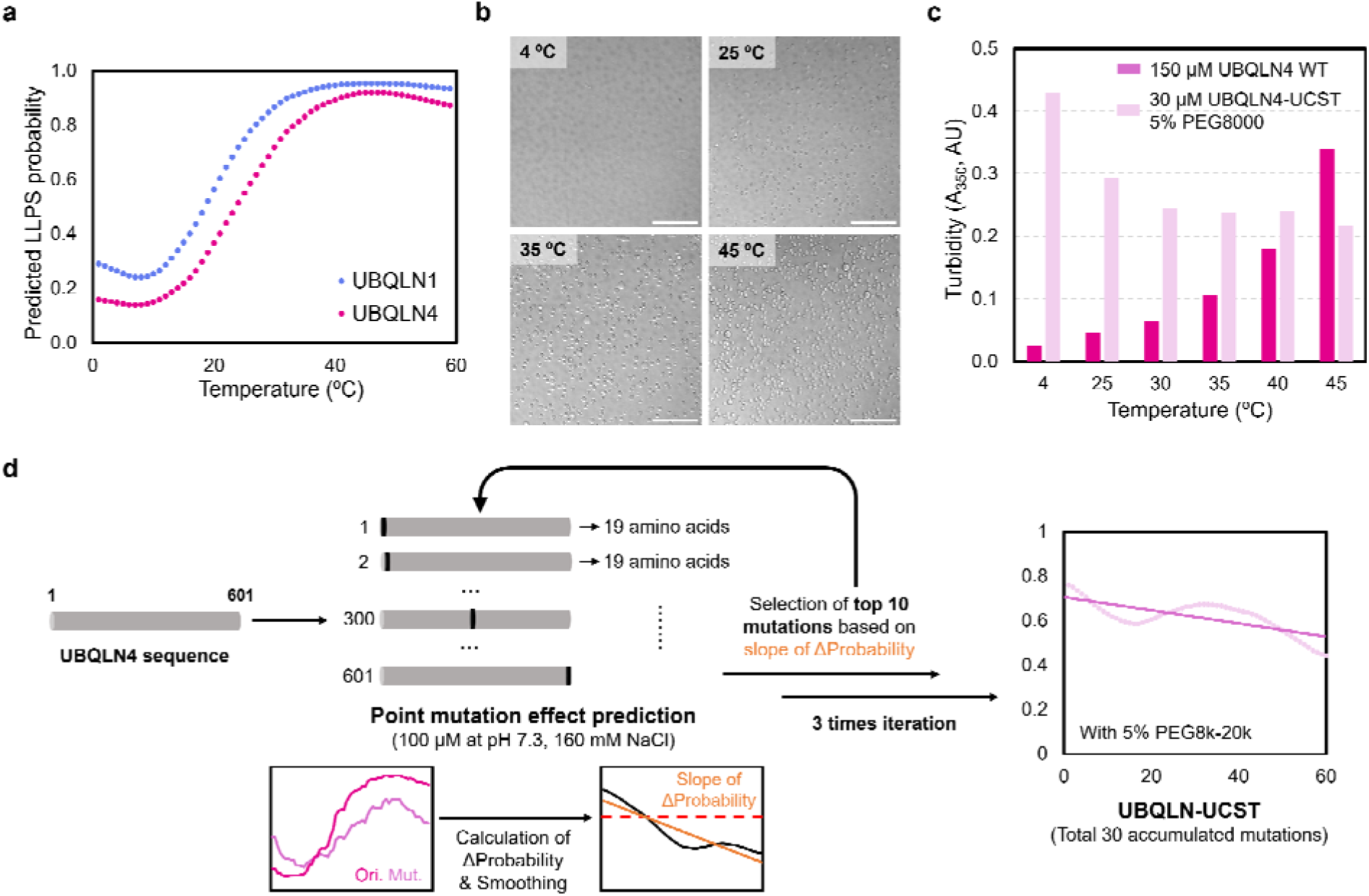
LLPSense-guided identification and rational reprogramming of UBQLN4 phase behavior. **(a)** Temperature-dependent LLPS probability profiles predicted by LLPSense for UBQLN1 and UBQLN4. The model accurately captures the LCST characteristics of both proteins. Predicted points were smoothed using a centered moving average (CMA) with a window size of 15. **(b)** Representative microscopy images of UBQLN4 WT condensates formed across a temperature gradient. Scale bars, 50 μm. **(c)** Experimental turbidity measurements (A_350_) comparing the thermal response of UBQLN4 WT and the UBQLN4-UCST variant. While the WT (dark magenta) exhibits distinct LCST behavior, the UBQLN4-UCST (light pink) displays a UCST-like profile. **(d)** Schematic workflow and computational strategy for the rational reprogramming of phase behavior. The design strategy prioritized the top *N=10* mutations in each round, selecting those with the largest ΔProbability in the desired direction (sign) to invert the condition-dependent profile. While the selection is based on individual single-mutation evaluations, the representative plot shows the simultaneous introduction of the top 10 mutations to better visualize the predicted shift in phase behavior.

To experimentally validate these predictions, we examined UBQLN4 *in vitro* and observed a clear increase in turbidity and condensate formation with increasing temperature, confirming its LCST nature (Fig. 5b, c). We next investigated whether LLPSense could guide the rational reprogramming of this LCST profile toward a UCST-like regime. Adapting a strategy analogous to the α-synuclein mutational screen, we prioritized mutations based on the slope of the difference in predicted LLPS probability between the variant and wild-type (ΔProbability) across the temperature gradient. We selected mutations with the most negative ΔProbability slopes, indicating a shift toward UCST behavior, and incorporated them into the sequence (Fig. 5d). To ensure a sufficient cumulative impact to invert the temperature dependence while safeguarding against loss of phase-separation propensity, mutations were introduced across three iterative rounds, with 10 substitutions accumulated per round (Supplementary Table 3). This stepwise engineering yielded a UBQLN4-UCST variant that formed condensates resembling percolated clusters (Supplementary Fig. 9b). The recombinantly expressed variant displayed the targeted inversion in temperature dependence, with markedly higher turbidity at lower temperatures than at elevated temperatures (Fig. 5c). These results demonstrate that LLPSense serves not only as a predictor of condition-dependent phase behavior but also as a tool for rationally tuning it through targeted sequence modifications.

### Proteome-wide analysis of physicochemical dependencies in phase separation

Having established the robustness of LLPSense, we applied the model to map the global landscape of protein phase separation across the human proteome under diverse physicochemical conditions. Increasing the concentration of macromolecular crowding agents (PEG 8k–20k) induced a proteome-wide shift toward higher predicted LLPS probabilities (Supplementary Fig. 10a). This trend aligns with the excluded volume effect, whereby macromolecular crowding increases the effective protein concentration and promotes condensation. Although these predicted scores may not perfectly correspond to absolute phase separation fractions, they provide a valuable framework for relative comparison, facilitating the benchmarking of the LLPS propensity of proteins of interest against the entire human proteome. On the other hand, increasing ionic strength by elevating NaCl concentrations generally reduced predicted LLPS probabilities (Supplementary Fig. 10c), indicating that electrostatic interactions contribute substantially to condensate formation for a large fraction of human proteins. Interestingly, increasing the NaCl concentration from 500 mM to 1000 mM resulted in a statistically significant increase in predicted probability (\*\*\*\**P* < 0.0001, Wilcoxon tests). This rebound suggests an increased contribution of hydrophobic interactions under ionic strengths far exceeding physiological levels.

We also analyzed the correlation between sequence characteristics and salt sensitivity, quantified by the slope of phase separation probability relative to salt concentration. We observed a negative correlation between this slope and the fraction of charged residues (FCR). This indicates that IDRs with higher overall charge density exhibit a steeper decline in phase separation stability due to electrostatic screening (Supplementary Fig. 11a–b). In contrast, because hydrophobic interactions are typically strengthened at higher ionic strengths, the hydropathy index exhibited a positive correlation with the salt-response slope (Supplementary Fig. 11c–d). Regarding net charge per residue (NCPR), we found that proteins with a near-neutral net charge (NCPR ≈ 0) show a higher prevalence of phase separation (Supplementary Fig. 11e). However, these proteins are also relatively sensitive to ionic screening (Supplementary Fig. 11f). Furthermore, it is theoretically expected that LLPS is energetically favored near the isoelectric point (pI), as the minimization of electrostatic repulsion facilitates protein–protein interactions. Accordingly, we analyzed the relationship between predicted scores and pH relative to pI (Supplementary Fig. 12). We found that basic proteins (high pI) sustained their phase separation probabilities even at pH levels exceeding 7.4, whereas acidic proteins (low pI) exhibited a decreasing trend in propensity as conditions became more alkaline. In summary, these results confirm that our model faithfully recapitulates the underlying physicochemical rules governing LLPS across diverse environmental landscapes at the proteome-wide scale.

## Discussion

In this study, we presented LLPSense, a unified machine learning framework that bridges intrinsic sequence determinants and extrinsic environmental factors to accurately predict protein phase separation. By harnessing high-dimensional embeddings from pre-trained protein language models, LLPSense overcomes the inherent limitations of previous sequence-only predictors, thereby enabling high-resolution characterization of phase behavior across diverse physicochemical landscapes.

Our comprehensive validation highlights the transformative potential of this condition-aware approach. First, LLPSense identified proteins exhibiting atypical phase-separation behavior, such as SGTA, in datasets previously curated as non-phase-separating. This finding reveals latent inaccuracies in current negative datasets and demonstrates the model’s ability to capture complex, reentrant phase behaviors. Secondly, by mapping the mutational landscape of α-synuclein, we demonstrated that the model can bidirectionally predict mutations that modulate phase separation propensity. This capability allows for the rapid identification of high-risk variants with pathogenic potential, effectively prioritizing candidates for experimental follow-up in LLPS-associated pathologies. Lastly, we successfully reprogrammed the thermodynamic profile of UBQLN4 from an LCST to a UCST regime through targeted sequence modification. This serves as a definitive proof-of-concept that LLPSense can facilitate the rational design of proteins with tunable, stimuli-responsive phase properties.

Furthermore, the streamlined modular architecture of LLPSense ensures future scalability. By leveraging embeddings as its core feature set, our framework is designed to adapt to the rapid advancements in deep learning. As more sophisticated protein language models (pLMs) emerge, LLPSense allows seamless integration of these next-generation representations. Concurrently, the predictive power of LLPSense is expected to grow as more high-quality experimental data are accumulated. This data-driven evolution will extend the framework’s applicability beyond binary classification, paving the way for more granular characterizations, such as distinguishing between diverse condensate material states. Building on the foundation of LLPSense, we plan to implement multifaceted strategies, including integrating structural features derived from AlphaFold 3, to develop models with enhanced predictive accuracy and broader generalizability.

By providing a reliable platform for predicting and manipulating condition-dependent phase separation, LLPSense enables rapid, proteome-wide analysis across diverse biological contexts. This versatile capability enables a broad range of investigations, from identifying disease-associated mutations to gaining deeper physical insight into cellular organization and the fundamental principles governing phase behavior. Additionally, this work lays the foundation for programmable biomolecular condensation, opening new avenues for innovation in synthetic biology and therapeutics development through synthetic condensates or by modulating LLPS.

## Methods

### Dataset preparation for LLPSeq training and evaluation

To ensure a fair and consistent comparison, LLPSeq was trained and evaluated using the dataset and evaluation strategy established by the PSPire model. Protein sequences for the PSPire dataset, identified by their UniProt accessions, were retrieved from the UniProt database (https://www.uniprot.org/) on March 29, 2024. While the positive protein sets (ID-PSPs and noID-PSPs) remained identical to those in the original study, a minor fraction of the negative set (non-PSPs) was excluded. This exclusion was necessary because subsequent UniProt sequence updates altered certain protein lengths, resulting in some exceeding the original PSPire filtering threshold (< 2,700 amino acids). Specifically, 30 out of 8,323 entries in the training set and 2 out of 1,961 entries in the testing set were removed. For the baseline comparison, 55 engineered features were extracted following the methodology described in the DeePhase study^7^. These features were used to train a baseline model to benchmark the performance of our protein language model (pLM)-based LLPSeq.

### Sampling of experimental condition data

Entries from LLPSDB v2 were processed into protein names, amino acid sequences, and condition-related variables, including protein concentration, temperature, pH, crowding agents, three types of salts (NaCl, KCl, and MgCl_), and glycerol percentage. Crowding agents were classified into six representative categories: PEG 300–1k, PEG 3k–6k, PEG 8k–20k, ficoll, dextran MW ≤ 40 kDa, and dextran MW ≥ 70 kDa. PEG and dextran with unspecified molecular weights were assigned to the PEG 3k–6k and Dextran MW ≤ 40 kDa categories, respectively. Experimental conditions were parsed from a structured Excel dataset into numerical feature vectors. Fields specified as fixed values or ranges (‘a-b’, ‘a-’, and ‘-b’) were interpreted according to predefined feature-specific limits, sampling densities, and sampling intervals (Supplementary Table 4). Closed ranges were uniformly sampled either as continuous intervals or as discrete unit-based grids, depending on the feature type. Open-ended ranges were expanded 10-fold above or 0.1-fold below the specified boundary, and subsequently clipped to the global minimum or maximum allowed for each feature. A Cartesian product of all condition values was generated to enumerate every valid experimental configuration. If the resulting combinatorial expansion exceeded a predefined limit (n > 100), a random subset of configurations was sampled. Finally, all features were normalized to the range [0, 1] using feature-specific maximum values.

### Model training and hyperparameter optimization

The LLPSense model was constructed using the eXtreme Gradient Boosting (XGBoost) framework, a scalable implementation of gradient-boosted decision trees. The model was trained with the ‘binary:logistic’ objective function to predict the probability of phase separation, minimizing the logarithmic loss (logloss) metric. To ensure robust generalization, we optimized key hyperparameters governing the tree architecture and learning process—specifically n_estimators, max_depth, learning_rate, min_child_weight, subsample, colsample_bytree, gamma, and regularization terms (reg_alpha, reg_lambda). Hyperparameters for both LLPSeq and LLPSense were optimized using the Optuna framework. For LLPSense, model selection was conducted using a 5-fold cluster-based cross-validation scheme. The final optimized hyperparameter configurations for each model are summarized in Supplementary Table 5.

### Standard prediction condition and phase-separating candidate screening

Unless otherwise specified, phase-separation probabilities were computed under standard conditions: 100 µM protein concentration, pH 7.3, 160 mM NaCl, and 25 °C. For candidate screening, we employed a temperature-based filtering approach, smoothing predicted probabilities with a centered moving-average (CMA) window of size 9. Candidates were identified as proteins exhibiting a probability exceeding 0.5 within the 10–50 °C range. For experimental validation, we selected SGTA alongside three randomly drawn proteins (ASCL1, FBLL1, and SEMG2) from a pool of sequences shorter than 1,000 amino acids, specifically those exhibiting a predicted phase transition threshold within the 10–50 °C screening window. The AlphaFold database structures^45^ of the proteins used in the experiments are shown in Supplementary Fig. 13.

### LLPS prediction of α-synuclein pathogenic mutations and comparison with catGRANULE

We curated LLPS modulation data for α-synuclein mutations from four independent studies. The gathered experimental evidence included mutational rankings derived from turbidity assays—specifically (a) E46K ≈ E83Q > A30G ≈ A53T > A30P ≥ WT > A53E > G51D^40^ and (b) E46K > H50Q ≈ A53T > WT ≈ A30P > A53E^41^— as well as (c) the identification of E46K, A53T, and H50Q as LLPS-enhancing mutations via droplet imaging^39, 42^. By consolidating these prior observations, we categorized the variants into four distinct groups relative to the wild-type (WT): significantly enhanced (E46K, E83Q), mildly enhanced (A30G, A53T, H50Q), comparable to WT (A30P), and suppressed (A53E, G51D). LLPSense was subsequently employed to predict the LLPS probability for each curated mutant (standard condition). For comparative benchmarking, we used the catGRANULE 2.0 ROBOT web server to generate mutation scores for the same set of variants, as well as for mutations identified at each iterative step of our study.

### Model-guided mutagenesis for design of UBQLN4-UCST

We employed an iterative model-guided mutagenesis strategy under standard prediction conditions in the absence of crowding agents. For each round, temperature-dependent probability curves were smoothed using a CMA with a window size of 15. The temperature sensitivity of each single-point mutant was quantified by calculating the slope of the difference in predicted LLPS probability between the variant and wild-type (ΔProbability = Probability_variant_−Probability_WT_) across the temperature gradient via linear regression. The comprehensive list of these mutations and their calculated ΔProbability slopes for each round is provided in Supplementary Table 3. We selected the top 10 mutations with the most negative ΔProbability slopes, indicating a shift from LCST to UCST behavior, and incorporated them into the sequence. This procedure was repeated for three consecutive rounds, resulting in a final UBQLN4-UCST variant harboring 30 cumulative mutations. Note that, for visualization in Fig. 5d, the probability curve for UBQLN4-UCST was plotted in the presence of 5% PEG 8k-20k, using a CMA window size of 15 for clarity. The AlphaFold database structure of UBQLN4 WT and the predicted structure^46^ of UBQLN4-UCST are shown in Supplementary Fig. 13. Finally, although these variants were initially identified using a version of LLPSense prior to its final refinement, the overall predicted temperature-dependent trends and selection logic remain consistent despite minor numerical discrepancies compared to the latest model version (Supplementary Fig. 14).

### Expression and purification of proteins

All proteins used in this study were cloned into the pET-21a(+) vector using the *NdeI* and *XhoI* restriction sites, allowing for expression with the vector-encoded C-terminal His-tag. Complete protein sequences are provided in Supplementary Table 6. The recombinant plasmids were transformed into *Escherichia coli* BL21(DE3) cells and cultured in LB medium supplemented with 100_µg/mL ampicillin at 37_°C. When the OD___ reached 0.4–0.7, protein overexpression was induced by adding 1 mM isopropyl β-D-1-thiogalactopyranoside (IPTG), followed by overnight incubation at 20 °C. Cells were harvested by centrifugation at 6,000_rpm (∼6,400 × g) and washed once with 1× PBS (160_mM NaCl, pH 7.3). The cell pellets were resuspended in Ni-IDA binding buffer (20 mM Tris-HCl, 150 mM NaCl, 10 mM imidazole, pH 8.0) and lysed by sonication. Lysates were clarified by centrifugation at 12,000_rpm (∼16,000 × g) for 15_min, and the supernatants were collected. The supernatant was loaded onto a Ni-IDA Excellose (Takara) resin column, washed with wash buffer (20 mM Tris-HCl, 150 mM NaCl, 50 mM imidazole, pH 8.0), and eluted with elution buffer (20 mM Tris-HCl, 150 mM NaCl, 300 mM imidazole, pH 8.0). For α-synuclein variants, the eluate was heat-shocked at 65 °C, then centrifuged at 13,000 rpm (∼16,000 × g) and filtered through a 0.2 µm syringe filter (ADVANTEC). Protein solutions were dialyzed against 1× PBS at 25 °C, then filtered through a 0.2 µm syringe filter. Protein concentrations were determined by measuring absorbance at 280 nm. Final protein solutions were aliquoted, flash-frozen in liquid nitrogen, and stored at −80_°C for long-term use.

### Turbidity assays

Protein phase separation was assessed in 96-well black plates with clear bottoms (Thermo Fisher). Turbidity was quantified by measuring absorbance at 350 nm using a spectral scanning multimode reader (VarioSkan Flash, Thermo Fisher). For each condition, 100 μL of protein solution was loaded per well. Measurements were recorded 30 min after the induction of phase separation, triggered by either the addition of a crowding agent or a temperature shift. Baseline correction was performed by subtracting the absorbance of PBS control wells containing the corresponding concentration of PEG8000. Each reported turbidity value represents the average of ten consecutive measurements taken from a single sample. In the specific case of α-synuclein variants, fifteen consecutive measurements were recorded per sample. Measurements were designed to identify consistent trends across diverse conditions for the same protein rather than repeated measurements under a single condition. All observed LLPS trends were independently replicated.

### Observation of protein condensates via microscopy

To visualize the protein condensates, differential interference contrast (DIC) imaging was performed using a laser-scanning confocal microscope (A1R HD25; Nikon) mounted on an Eclipse Ti2 inverted microscope, equipped with a 60×/1.40 numerical aperture (NA) apochromatic oil-immersion objective. To minimize non-specific surface adsorption, all imaging experiments were performed using BSA-passivated 96-well plates. Surface passivation was conducted by incubating the wells with 0.25% (w/v) BSA in PBS for 2 h at room temperature, followed by twice-washing with PBS to remove residual BSA. Prior to image acquisition, 150 µL of protein samples were loaded into passivated wells and allowed to settle for 60 min (unless otherwise specified) to ensure droplet formation and equilibration.

### Proteome retrieval and physicochemical profiling

We obtained the Swiss-Prot reviewed human proteome from the UniProt database (accessed on November 25, 2025). For the analysis, we specifically selected proteins containing fewer than 4,000 residues. Sequence-based physicochemical properties, including the fraction of charged residues (FCR), net charge per residue (NCPR), and hydropathy (0-9 scale), were calculated using the sparrow library (https://sparrow-online.com).

### Implementation details

All computational experiments were conducted using a single NVIDIA RTX A6000 GPU. Feature extraction utilizing the ProtT5 model required approximately 131 ms per protein on average (benchmarked on the human proteome restricted to lengths under 4,000 residues). For inference, LLPSense processed a single protein-condition pair in approximately 34 ms. Notably, inference consumed less than 8 GB of VRAM, demonstrating the framework’s computational efficiency for large-scale screening.

## Data availability

The pre-established models, curated dataset, and input template file are publicly available on GitHub at https://github.com/NearNiah/LLPSense. The dataset was curated based on LLPSDB v2.0 (http://bio-comp.org.cn/llpsdb). Source data are provided with this paper.

## Code availability

The codes of LLPSense and LLPSeq developed in this study are available at https://github.com/NearNiah/LLPSense.

## Acknowledgements

This work was supported by the National Research Founda-tion of Korea (NRF) grants (2023R1A2C2005183) and the Global Science Research Center Program (RS-2024-00411134) funded by the Korean government (MSIT).

## Author contributions

J.B. and M.K. conceived the study, developed the model, and wrote the code. D.L. curated the datasets under the guidance of J.B. J.B. performed the experiments with support from D.L. M.K. performed the computational analysis and evaluated the model performance. Y.J. provided guidance on the experimental design and data interpretation. J.B. conducted the data analysis, visualized the results, and drafted the manuscript. Y.J. and M.K. reviewed and edited the manuscript. K.-J.Y. and Y.J. supervised the project.

