## Supplementary Information for "A machine learning framework for predicting and modulating condition-dependent protein phase separation"

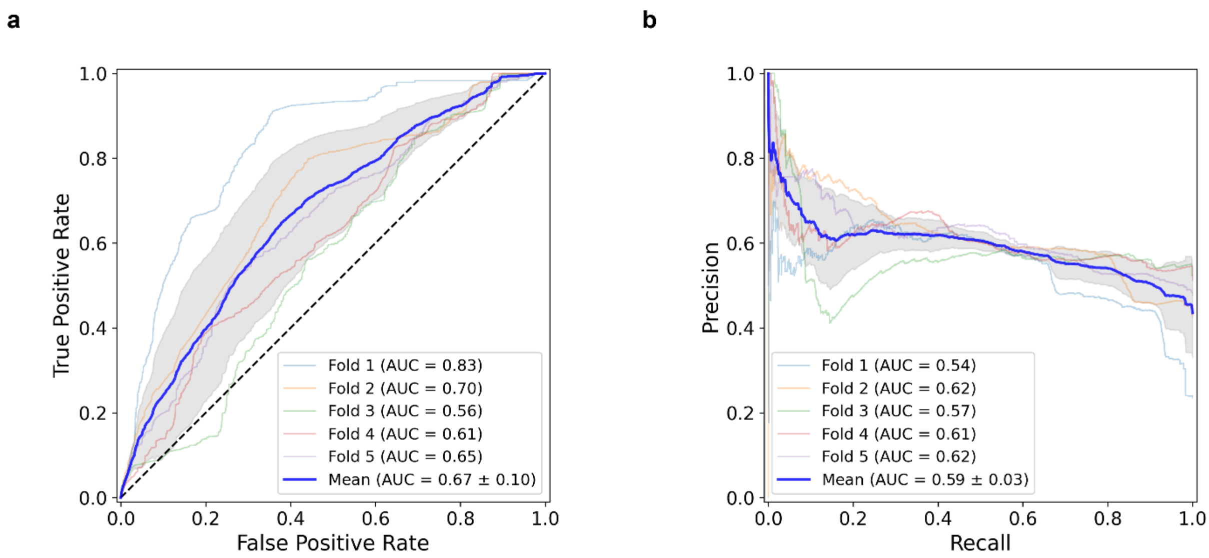


**Supplementary Figure 1. Performance of Droppler retrained on our expanded dataset using a cluster-based 5-fold cross-validation. (a)** Receiver operating characteristic (ROC) curves. **(b)** Precision-recall (PR) curves. Gray area, ± 1 standard deviations.

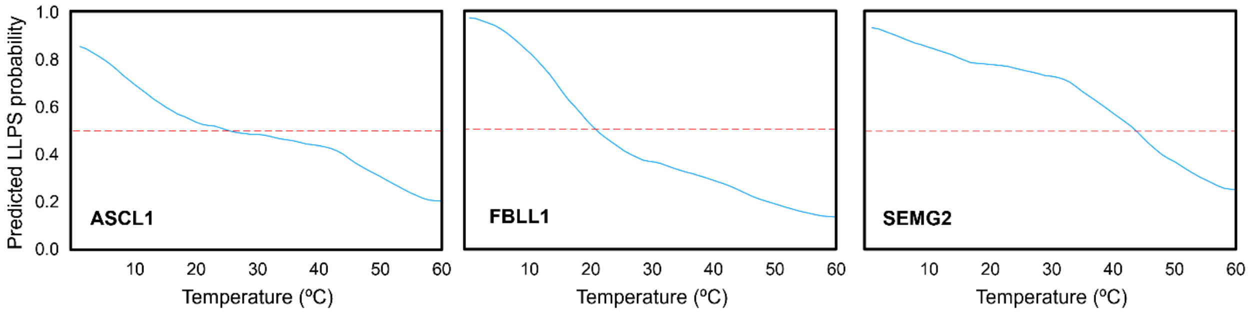


**Supplementary Figure 2. LLPSense prediction plot for the randomly selected ASCL1, FBLL1, and SEMG2 from the list of candidates for the *in vitro* experiment.** The data was smoothed using a centered moving average (window size = 15).

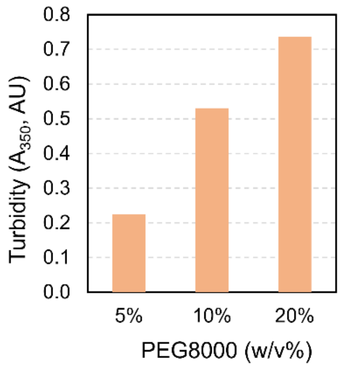


**Supplementary Figure 3. Turbidity measurements ($A_{350}$) for 12.5 µM SEMG2 under crowding conditions.**

$A_{350}$ of wild type (WT)

| WT | 80 µM | 40 µM | 20 µM | 10 µM |
|---|---|---|---|---|
| 20% PEG | 0.279 | 0.154 | 0.085 | 0.061 |
| 10% PEG | 0.099 | 0.090 | 0.068 | |
| 5% PEG | 0.007 | 0.055 | | |

≥ 0.250
0.200
0.175
0.150
0.125
0.100
0.075
0.050
0.025
≤ 0.010
Turbidity (AU)

ΔTurbidity = $A_{350}$ of variant - $A_{350}$ of wild type

| E1 | 80 µM | 40 µM | 20 µM | 10 µM |
|---|---|---|---|---|
| 20% PEG | 0.049 | 0.021 | 0.023 | 0.005 |
| 10% PEG | -0.013 | -0.005 | 0.001 | |
| 5% PEG | 0.030 | 0.013 | | |

| E2 | 80 µM | 40 µM | 20 µM | 10 µM |
|---|---|---|---|---|
| 20% PEG | 0.119 | 0.056 | 0.039 | 0.021 |
| 10% PEG | 0.100 | 0.033 | 0.007 | |
| 5% PEG | 0.078 | 0.030 | | |

LLPS-enhanced variants

| S1 | 80 µM | 40 µM | 20 µM | 10 µM |
|---|---|---|---|---|
| 20% PEG | -0.078 | -0.020 | -0.003 | -0.001 |
| 10% PEG | -0.020 | -0.008 | -0.002 | |
| 5% PEG | 0.009 | 0.003 | | |

| S2 | 80 µM | 40 µM | 20 µM | 10 µM |
|---|---|---|---|---|
| 20% PEG | -0.072 | -0.025 | -0.009 | 0.003 |
| 10% PEG | -0.069 | -0.038 | -0.016 | |
| 5% PEG | -0.002 | 0.003 | | |

LLPS-suppressed variants

| E28P | 80 µM | 40 µM | 20 µM | 10 µM |
|---|---|---|---|---|
| 20% PEG | -0.036 | 0.003 | 0.012 | 0.003 |
| 10% PEG | -0.027 | -0.013 | -0.008 | |
| 5% PEG | 0.021 | 0.010 | | |

| E130M | 80 µM | 40 µM | 20 µM | 10 µM |
|---|---|---|---|---|
| 20% PEG | -0.001 | -0.002 | 0.018 | 0.005 |
| 10% PEG | -0.066 | -0.029 | -0.016 | |
| 5% PEG | 0.001 | 0.005 | | |

Control variants (vs. E1)

≥ 0.050
0.025
0.000
-0.025
≤ -0.050
ΔTurbidity

**Supplementary Figure 4. Heatmaps of turbidity of α-synuclein WT and ΔTurbidity of each variant.** ΔTurbidity was calculated by subtracting the WT turbidity from that of each variant under identical experimental conditions. Variants are categorized into LLPS-enhanced (E1, E2), LLPS-suppressed (S1, S2), and control (E28P, E130M) groups.

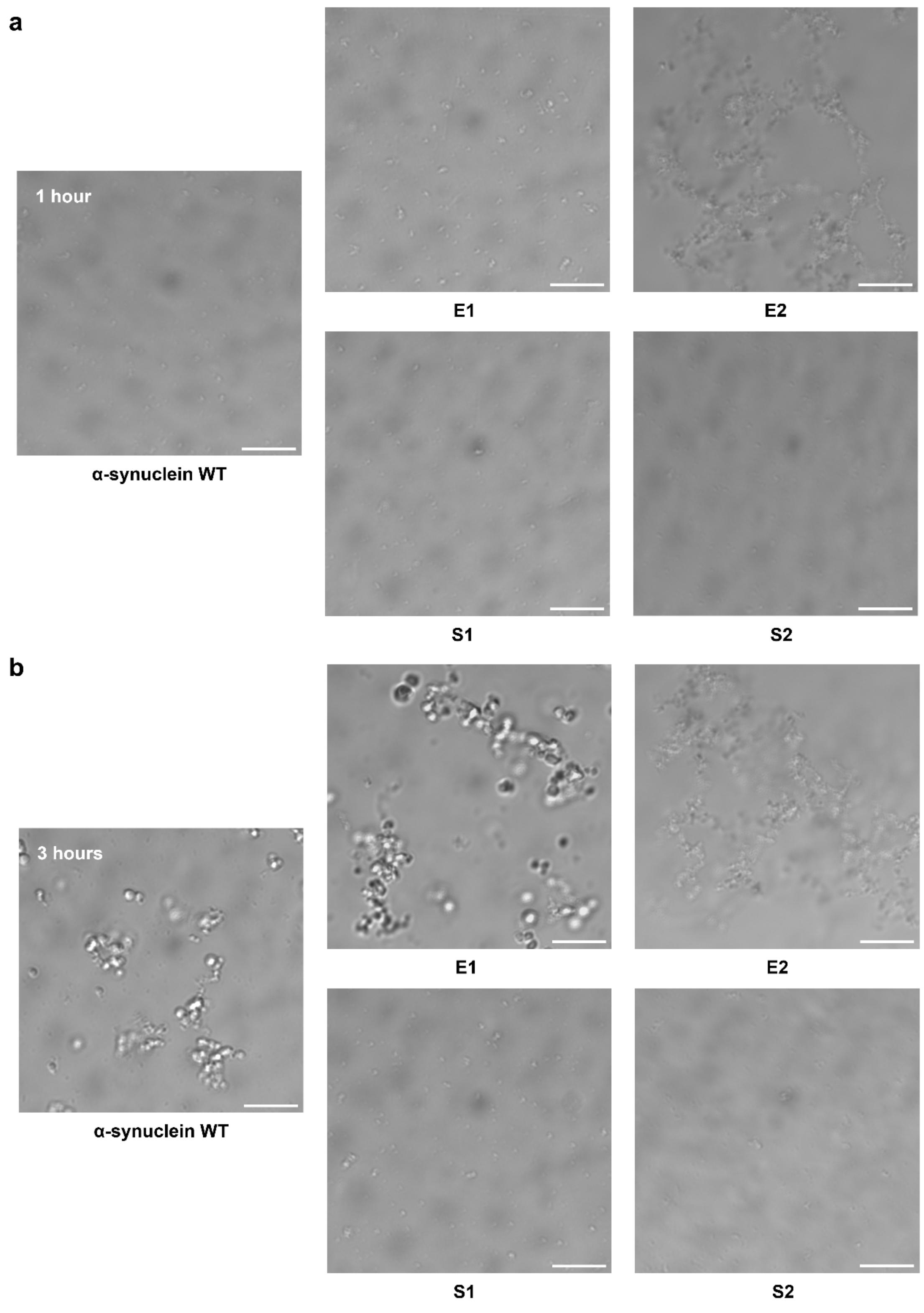


**Supplementary Figure 5. Time-dependent formation of α-synuclein condensates under crowded conditions.** Representative differential interference contrast (DIC) microscopic images showing protein condensates formed by α-synuclein wild-type (WT) and its variants (E1, E2, S1, and S2) at different time points. All samples were imaged in PBS buffer containing 100 μM protein and 20% (w/v) PEG-8000 to induce phase separation. The panels illustrate the presence and relative abundance of condensates for each protein variant (a) 1 hour and (b) 3 hours after sample preparation under identical experimental conditions. Scale bars, 10 μm.

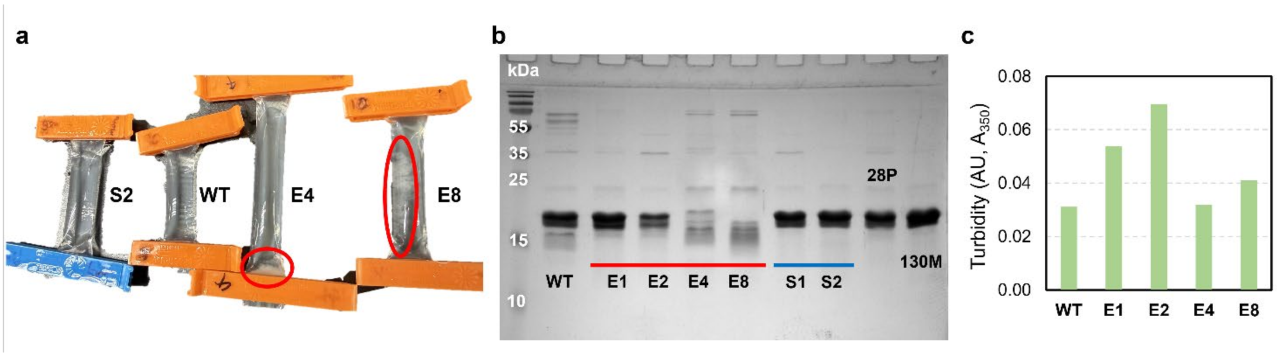


**Supplementary Figure 6. Aggregation and instability of higher-order enhanced α-synuclein mutants. (a)** Pronounced protein aggregation (white precipitate, red circles) in the dialysis cassettes of E4 and E8 after PBS dialysis. **(b)** SDS-PAGE analysis (15% polyacrylamide denaturing gel) of all purified α-synuclein variants. The E4 and E8 lanes show significantly reduced protein concentration in the main band and exhibit degradation smears. **(c)** Turbidity measurements ($A_{350}$) of protein variants in the presence of 20% PEG8000 and 20 μM protein. E1 and E2 exhibit the highest turbidity levels. Notably, E4 and E8 show higher turbidity compared to WT; despite their lower purity profiles observed in (b), these results suggest an increased propensity for phase transition.

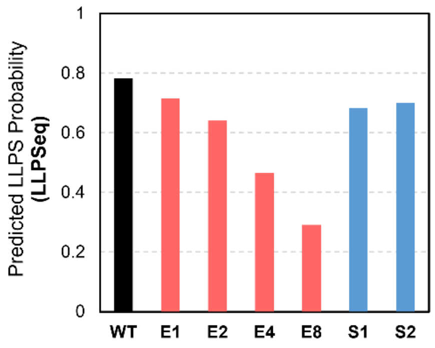


**Supplementary Figure 7. Predicted probabilities by LLPSeq for LLPSense-selecte variants.** Despite being selected for enhanced phase separation, these variants (E1–E8) show a monotonic decrease in probability according to LLPSeq, further highlighting LLPSeq's inability to resolve gain-of-function sequence alterations.

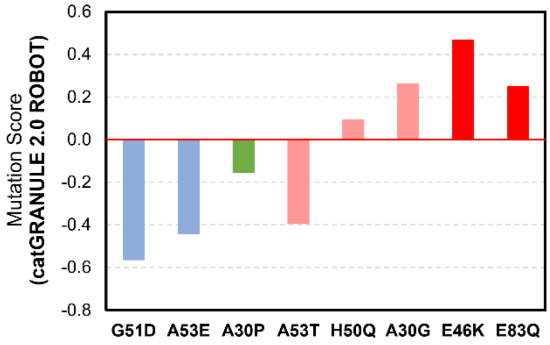


**Supplementary Figure 8. Prediction of α-synuclein mutation scores using catGRANULE 2.0 ROBOT.** The bar graph shows the mutation scores for various α-synuclein variants predicted by the catGRANULE 2.0 ROBOT web server. The x-axis represents specific point mutations, and the y-axis indicates the predicted mutation score of each mutations. The variants are color-coded according to their experimental liquid-liquid phase separation (LLPS) propensity reported in previous literature[1-4]: red (significantly enhanced), pink (mildly enhanced), green (comparable), and blue (suppressed) compared to WT. This comparison highlights the performance and limitations of the conventional prediction tool in capturing the experimentally observed LLPS behavior of α-synuclein variants.

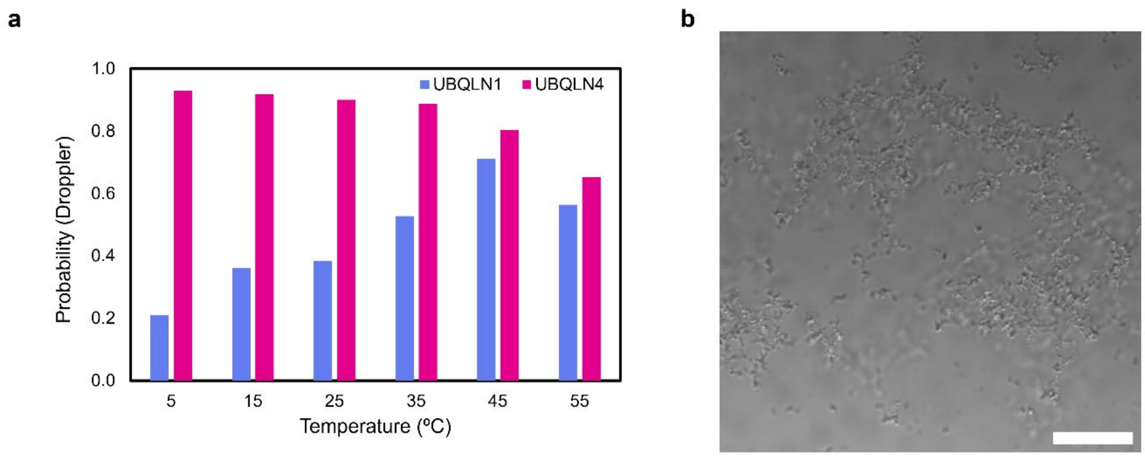


**Supplementary Figure 9. (a)** Temperature-dependent LLPS probability predictions of UBQLN family from the Droppler model (retrained on our expanded dataset). **(b)** Representative microscopy image of the LLPSense-designed UBQLN4-UCST mutant. Scale bar, 50 µm.

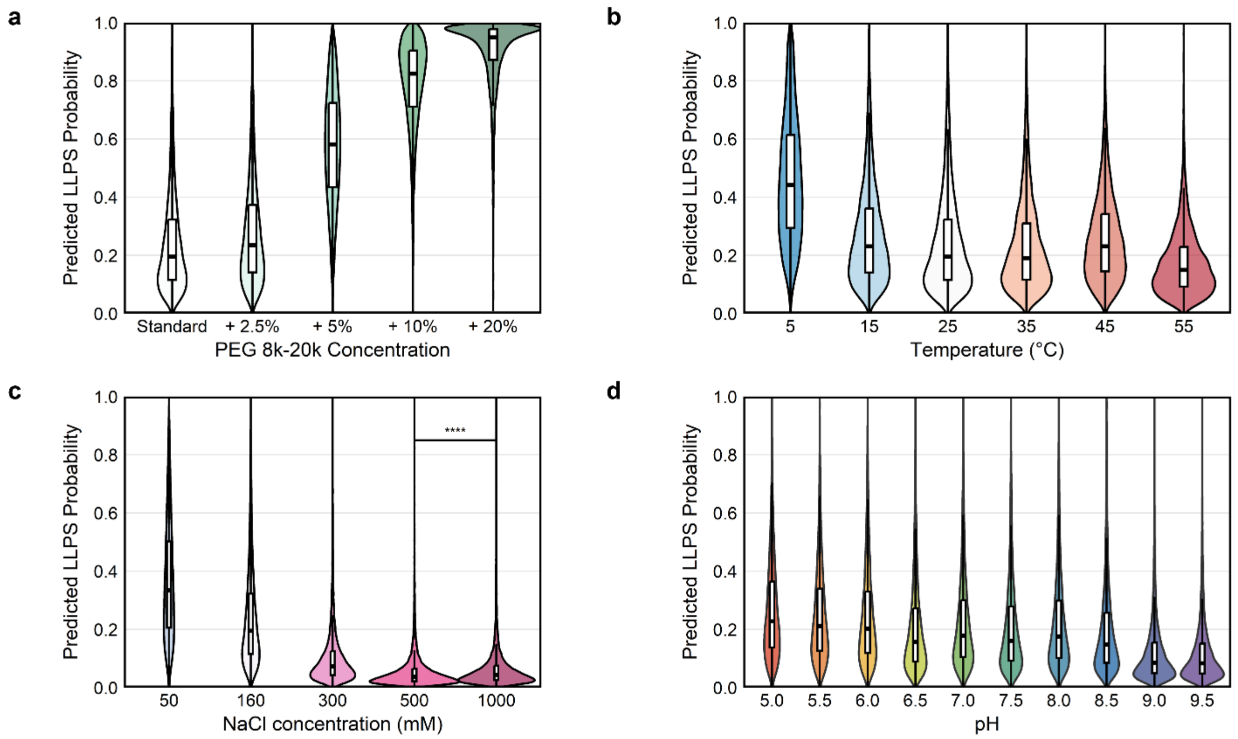


**Supplementary Figure 10. Distribution of predicted LLPS probabilities in the human proteome under various physicochemical conditions.** Violin plots illustrate the LLPSense-predicted phase separation probability for the entire human proteome across different environmental parameters. **(a)** Effect of macromolecular crowding. LLPS probability distribution as a function of PEG 8k-20k concentration (ranging from 0% [Standard] to 20%). Increasing concentration of the crowding agent significantly shifts the proteome towards higher LLPS probabilities, consistent with the excluded volume effect. **(b)** Temperature dependence. Predicted LLPS probabilities at temperatures ranging from 5 °C to 55 °C. **(c)** Influence of ionic strength. LLPS probabilities across various NaCl concentrations (50 mM to 1000 mM). Higher salt concentrations generally lead to a decrease in predicted LLPS probability, suggesting the dominance of electrostatic interactions in the formation of these condensates. For the comparison between 500 mM and 1000 mM, statistical significance was confirmed (****$P < 0.0001$ using paired one-sided [greater], paired two-sided, and two-sided rank-sum Wilcoxon tests). **(d)** pH dependence. Distribution of LLPS probabilities across a pH range from 5.0 to 9.5. Standard condition definition: All simulations, unless otherwise specified, were performed under standard physiological conditions: pH 7.3, 160 mM NaCl, 25 °C, and in the absence of crowding agents.

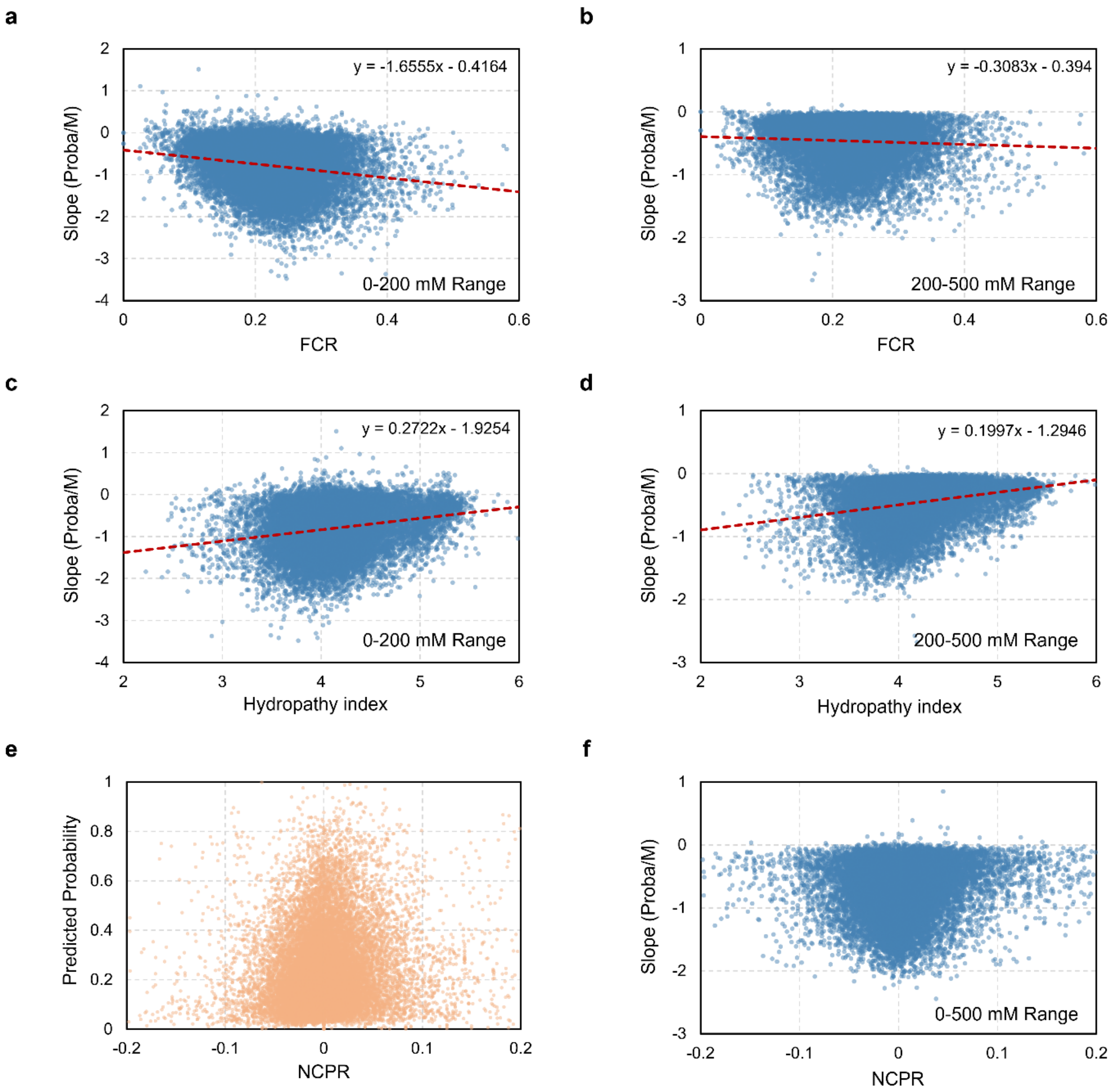


**Supplementary Figure 11. Correlation between sequence characteristics and salt sensitivity of phase separation. (a–b)** Correlation between the fraction of charged residues (FCR) and the salt-response slope (change in phase separation probability per molar concentration of salt) in the (a) 0–200 mM and (b) 200–500 mM salt concentration ranges. The red dashed lines represent linear regression fits, showing a negative correlation indicative of electrostatic screening. **(c–d)** Correlation between the hydropathy index (normalized to a 0–9 scale based on Kyte-Doolittle values) and the salt-response slope in the (c) 0–200 mM and (d) 200–500 mM ranges. The positive correlation suggests that hydrophobic interactions are enhanced at higher ionic strengths. **(e)** Predicted phase separation probability as a function of net charge per residue (NCPR), showing that LLPS prevalence peaks near NCPR ≈ 0. **(f)** Relationship between NCPR and the salt-response slope calculated across the 0–500 mM salt range. For all panels, axes are zoomed into the major distribution area of the data points.

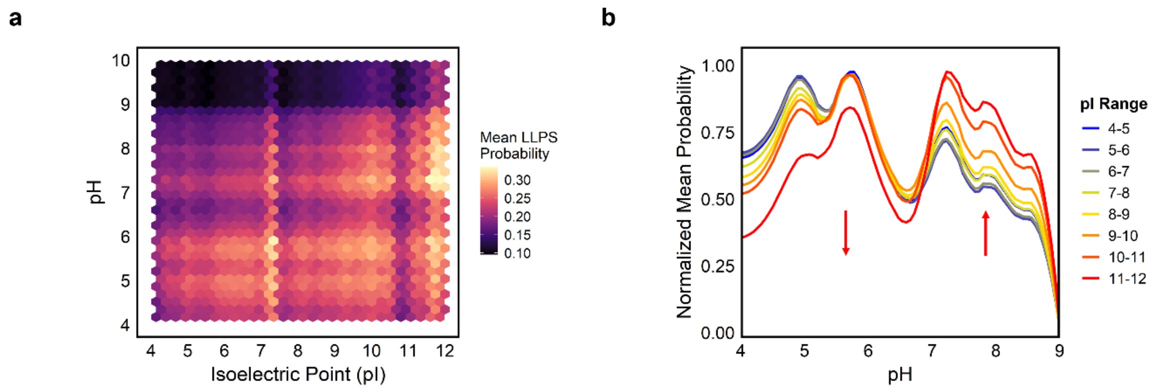


**Supplementary Figure 12. Dependence of phase separation probability on solution pH and protein isoelectric point (pI). (a)** Hexbin plot showing the mean predicted LLPS probability as a function of the isoelectric point (pI) and solution pH. The color scale indicates the mean probability. **(b)** Normalized mean LLPS probability across a pH range of 4–9, categorized by protein pI ranges. The trend lines were generated using LOESS (Locally Estimated Scatterplot Smoothing) with a span of 0.2. The red arrows indicate the shifting trends in phase separation propensity as pI increases.

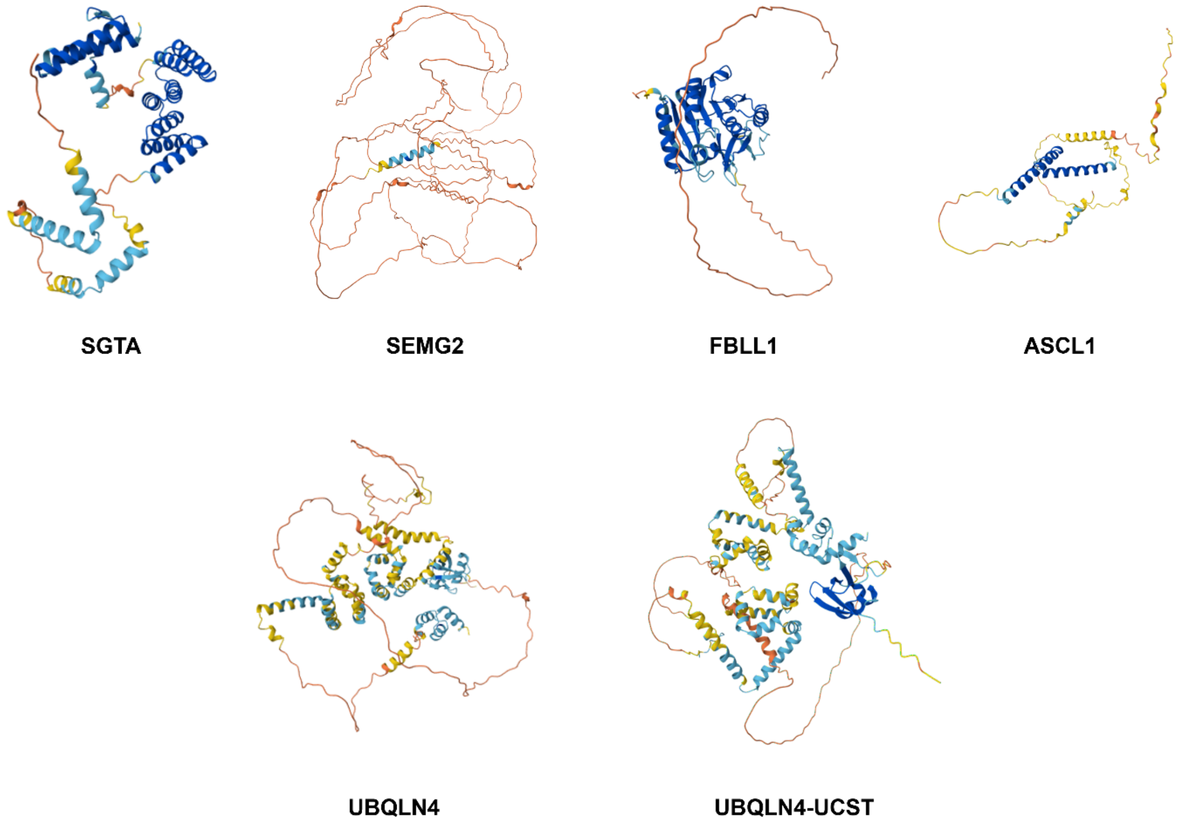


**Supplementary Figure 13. AlphaFold-predicted structures of proteins used in this study.** The structures for SGTA, SEMG2, FBLL1, ASCL1, and wild-type (WT) UBQLN4 were retrieved from the AlphaFold database[5]. The structure for the UBQLN4-UCST variant was predicted using the AlphaFold web server[6].

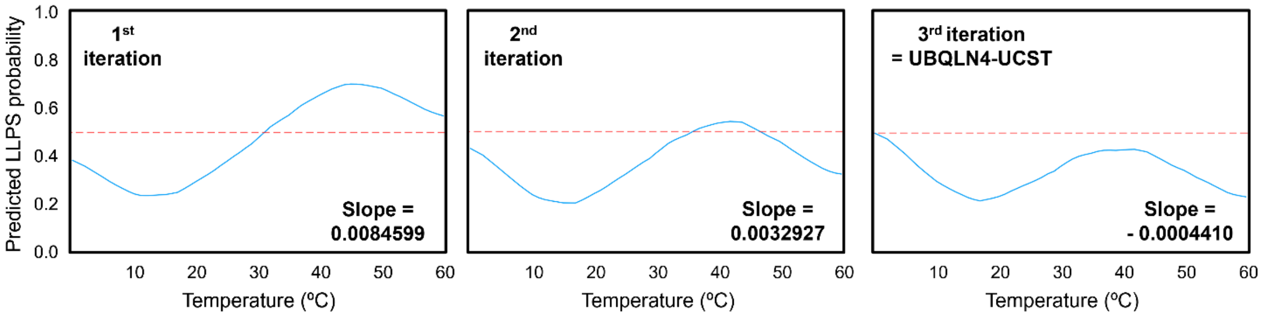


**Supplementary Figure 14. Iterative transition of predicted LLPS probability profiles for UBQLN4.** The panels illustrate the step-by-step conversion of UBQLN4 from an LCST-type to a UCST-type protein across three rounds of model-guided mutagenesis. These probability curves and their corresponding slope values (probability/°C) were generated using the final version of LLPSense. The progressive shift from a positive slope to a negative slope demonstrates the successful computational reprogramming of the temperature-dependent phase behavior. A complete list of the mutations for each round is provided in Supplementary Table 2.

**Supplementary Table 1. List of LLPSense-designed α-synuclein variants and their predicted phase separation probabilities.** The 'E' and 'S' prefixes denote variants designed to exhibit enhanced and suppressed phase separation propensity, respectively. The numerical suffix indicates the cumulative number of mutations relative to the wild-type (WT) sequence. Mutations were introduced iteratively; for example, higher-order variants (e.g., E2, E4) retain the substitutions from the preceding variant while incorporating additional mutations. Predicted LLPS probability values were computed by LLPSense under standard conditions (100 µM protein concentration, 160 mM NaCl, pH 7.3, at 25 °C).

| Variant name | Added mutation(s) | Predicted LLPS probability at standard condition |
|---|---|---|
| E1 | WT + E28M | 0.257 |
| E2 | E1 + E57F | 0.347 |
| E4 | E2 + A140W + E20R | 0.442 |
| E8 | E4 + A76R + E35Q + E105R + V118M | 0.607 |
| S1 | WT + V77D | 0.091 |
| S2 | S1 + M127E | 0.068 |

**Supplementary Table 2. catGRANULE 2.0 mutation scores for α-synuclein mutations identified by LLPSense across iterative steps.** Predicted mutation scores were calculated using the catGRANULE 2.0 ROBOT web server. Each row represents a mutation identified during the iterative optimization steps of LLPSense, progressing from wild-type (WT) to successive stages (E1–E8 and S1–S2).

| Mutation step | Mutation | Predicted mutation score with catGRANULE 2.0 ROBOT |
|---|---|---|
| WT to E1 | E28M | -0.0481 |
| E1 to E2 | E57F | 0.024057 |
| E2 to E3 | A140W | -0.55322 |
| E3 to E4 | E20R | 0.652005 |
| E4 to E5 | A76R | 0.204309 |
| E5 to E6 | E35Q | 0.252014 |
| E6 to E7 | E105R | 0.646873 |
| E7 to E8 | V118M | 0.107316 |
| WT to S1 | V77D | 0.30061 |
| S1 to S2 | M127E | 0.047995 |
| WT to Residue-specific control | E28P | 0.288585 |
| WT to Position-specific control | E130M | -0.0481 |

**Supplementary Table 3. LLPSense-guided mutagenesis of UBQLN4 and the magnitude of predicted slope of ΔProbability for each mutation.** Mutations were selected to invert the temperature dependence of UBQLN4 from an LCST to a UCST regime. The mutagenesis was performed over three iterative rounds, with ten single-point substitutions introduced per round. Values listed for each mutation represent the slope (probability/°C) of the difference in predicted LLPS probability between the variant and wild-type (ΔProbability = $\text{Probability}_{\text{variant}} - \text{Probability}_{\text{WT}}$).

| | | | | | | | | | | |
|---|---|---|---|---|---|---|---|---|---|---|
| 1st iteration | M200C | L272C | M569C | M229C | P394C | M218C | M451C | I219C | M225Y | F378C |
| | -0.0060 | -0.0059 | -0.0055 | -0.0054 | -0.0053 | -0.0051 | -0.0050 | -0.0049 | -0.0048 | -0.0048 |
| 2nd iteration | R31I | Y405K | I60D | V78Y | M537K | L460D | P12N | R10N | L76E | F306D |
| | -0.0023 | -0.0020 | -0.0018 | -0.0018 | -0.0018 | -0.0018 | -0.0018 | -0.0017 | -0.0017 | -0.0017 |
| 3rd iteration | A591D | I199P | L67D | F571P | P203L | L61D | I582P | L282D | T248P | M285D |
| | -0.0013 | -0.0013 | -0.0013 | -0.0013 | -0.0013 | -0.0012 | -0.0012 | -0.0012 | -0.0012 | -0.0012 |

**Supplementary Table 4. Predefined feature-specific limits, sampling densities, and intervals.**

| Conditions | Number of sampling | Minimum value | Maximum value | Sampling interval |
|---|---|---|---|---|
| **Temperature (°C)** | 50 | 0 | 60 | 1 |
| **Protein concentration** | 10 | 0 | 1000 | continuous |
| **pH** | 10 | 0 | 14 | 0.1 |
| **[$MgCl_2$] (mM)** | 10 | 0 | 50 | 1 |
| **[NaCl] (mM)** | 50 | 0 | 2000 | 10 |
| **[KCl] (mM)** | 50 | 0 | 1000 | 10 |
| **Crowding agents (6 categories, %)** | 30 | 0 | 50 | 1 |
| **Glycerol (%)** | 10 | 0 | 10 | 1 |

**Supplementary Table 5. Optimized hyperparameters.**

| Hyperparameters | LLPSeq | LLPSense |
|---|---|---|
| **n_estimators** | 956 | 877 |
| **max_depth** | 14 | 14 |
| **learning_rate** | 0.022836067655687835 | 0.023722174554030036 |
| **min_child_weight** | 1 | 2 |
| **colsample_bytree** | 0.8073843951528245 | 0.5655509178160505 |
| **subsample** | 0.7120210675891471 | 0.5482175662874387 |
| **gamma** | 1.3626081386701552e-05 | 0.0012682360748828933 |
| **reg_lambda** | 72.93174736264936 | 0.3024802937837069 |
| **reg_alpha** | 0.2636053564337023 | 0.04238171400820899 |

**Supplementary Table 6. Protein sequences for *in vitro* experiments.** The amino acid sequences enclosed in parentheses represent the C-terminal His-tag originating from the pET-21a(+) vector. Mutated residues in the UBQLN4-UCST sequence are highlighted in **bold**.

| Protein names | Amino acid sequences |
|---|---|
| **SGTA** | MDNKKRLAYAIIQFLHDQLRHGGLSSDAQESLEVAIQCLETAFGVTVEDSDLALPQTLPE<br>IFEAAATGKEMPQDLRSPARTPPSEEDSAEAERLKTEGNEQMKVENFEAAVHFYGKAIE<br>LNPANAVYFCNRAAAYSKLGNYAGAVQDCERAICIDPAYSKAYGRMGLALSSLNKHVE<br>AVAYYKKALELDPDNETYKSNLKIAELKLREAPSPTGGVGSFDIAGLLNNPGFMSMASN<br>LMNNPQIQQLMSGMISGGNNPLGTPGTSPSQNDLASLIQAGQQFAQQMQQQNPELIEQL<br>RSQIRSRTPSASNDDQQE(LEHHHHHH) |
| **ASCL1** | MESSAKMESGGAGQQPQPQPQQPFLPPAACFFATAAAAAAAAAAAAAQSAQQQQQQ<br>QQQQQAPQLRPAADGQPSGGGHKSAPKQVKRQRSSSPELMRCKRRLNFSGFGYSLPQQ<br>QPAAVARRNERERNRVKLVNLGFATLREHVPNGAANKKMSKVETLRSAVEYIRALQQL<br>LDEHDAVSAAFQAGVLSPTISPNYSNDLNSMAGSPVSSYSSDEGSYDPLSPEEQELLDFT<br>NWF(LEHHHHHH) |
| **FBLL1** | MKSAASSRGGGGGGRGGGGWGSWGGGRGGGGGAGKGGGGDGGGQGGKGGFGARA<br>RGFGGGGRGRGRGGGDGKDRGGGGQRRGGVAKSKSRRRKGAMVVSVEPHRHEGVFI<br>YRGAEDALVTLNMVPGQSVYGERRVTVTEGGVKQEYRTWNPFRSKLAAAILGGVDQI<br>HIKPKSKVLYLGAASGTTVSHVSDIIGPDGLVYAVEFSHRAGRDLVNVAKKRTNIIPVLE<br>DARHPLKYRMLIGMVDVIFADVAQPDQSRIVALNAHTFLRNGGHFLISIKANCIDSTASA<br>EAVFASEVRKLQQENLKPQEQLTLEPYERDHAVVVGVYRPLPKSSSK(LEHHHHHH) |
| **SEMG2** | MKSIILFVLSLLLILEKQAAVMGQKGGSKGQLPSGSSQFPHGQKGQHYFGQKDQQHTK<br>SKGSFSIQHTYHVDINDHDWTRKSQQYDLNALHKATKSKQHLGGSQQLLNYKQEGRD<br>HDKSKGHFHMIVIHHKGGQAHHGTQNPSQDQGNSPSGKGLSSQCSNTEKRLWVHGLS<br>KEQASASGAQKGRTQGGSQSSYVLQTEELVVNKQQRETKNSHQNKGHYQNVVDVRE<br>EHSSKLQTSLHPAHQDRLQHGPKDIFTTQDELLVYNKNQHQTKNLSQDQEHGRKAHKI<br>SYPSSRTEERQLHHGEKSVQKDVSKGSISIQTEEKIHGKSQNQVTIHSQDQEHGHKENKI<br>SYQSSSTEERHLNCGEKGIQKGVSKGSISIQTEEQIHGKSQNQVRIPSQAQEYGHKENKI<br>SYQSSSTEERRLNSGEKDVQKGVSKGSISIQTEEKIHGKSQNQVTIPSQDQEHGHKENK<br>MSYQSSSTEERRLNYGGKSTQKDVSQSSISFQIEKLVEGKSQIQTPNPNQDQWSGQNAK<br>GKSGQSADSKQDLLSHEQKGRYKQESSESHNIVITEHEVAQDDHLTQQYNEDRNPIST(L<br>EHHHHHH) |
| **α-synuclein (WT)** | MDVFMKGLSKAKEGVVAAAEKTKQGVAEAAGKTKEGVLYVGSKTKEGVVHGVATVA<br>EKTKEQVTNVGGAVVTGVTAVAQKTVEGAGSIAAATGFVKKDQLGKNEEGAPQEGILE<br>DMPVDPDNEAYEMPSEEGYQDYEPEA(LEHHHHHH) |
| **UBQLN4 (WT)** | MAEPSGAETRPPIRVTVKTPKDKEEIVICDRASVKEFKEEISRRFKAQQDQLVLIFAGKIL<br>KDGDTLNQHGIKDGLTVHLVIKTPQKAQDPAAATASSPSTPDPASAPSTTPASPATPAQPS<br>TSGSASSDAGSGSRRSSGGGPSPGAGEGSPSATASILSGFGGILGLGSLGLGSANFMELQ<br>QQMQRQLMSNPEMLSQIMENPLVQDMMSNPDLMRHMIMANPQMQQLMERNPEISHM<br>LNNPELMRQTMELARNPAMMQEMMRNQDRALSNLESIPGGYNALRRMYTDIQEPMFS<br>AAREQFGNNPFSSLAGNSDSSSSQPLRTENREPLPNPWSPSPPTSQAPGSGGEGTGGSGT<br>SQVHPTVSNPFGINAASLGSGMFNSPEMQALLQQISENPQLMQNVISAPYMRSMMQTL<br>AQNPDFAAQMMVNVPLFAGNPQLQEQLRLQLPVFLQQMQNPESLSILTNPRAMQALLQ<br>IQQGLQTLQTEAPGLVPSLGSFGISRTPAPSAGSNAGSTPEAPTSSPATPATSSPTGASSAQ<br>QQLMQQMIQLLAGSGNSQVQTPEVRFQQQLEQLNSMGFINREANLQALIATGGDINAAI<br>ERLLGSQLS(LEHHHHHH) |
| **UBQLN4-UCST** | MAEPSGAET**N**P**N**IRVTVKTPKDKEEIVICD**I**ASVKEFKEEISRRFKAQQDQLVLIFAGK**D**<br>**D**KDGDT**D**NQHGIKDG**E**T**Y**HLVIKTPQKAQDPAAATASSPSTPDPASAPSTTPASPATPAQ<br>PSTSGSASSDAGSGSRRSSGGGPSPGAGEGSPSATASILSGFGGILGLGSLGLGSANFMEL<br>QQQMQRQLMSNPEMLSQ**PC**EN**L**LVQDMMSNPDLMRH**CC**MANPQ**Y**QQL**C**ERNPEISH<br>MLNNPELMRQ**P**MELARNPAMMQEMMRNQDRALSN**C**ESIPGGYNA**D**RR**D**YTDIQEPM<br>FSAAREQFGNNP**D**SSLAGNSDSSSSQPLRTENREPLPNPWSPSPPTSQAPGSGGEGTGGS<br>GTSQVHPTVSNPFGINAASLGSGM**C**NSPEMQALLQQISEN**C**QLMQNVISAP**K**MRSMM<br>QTLAQNPDFAAQMMVNVPLFAGNPQLQEQLRLQLPVFLQQ**C**QNPESLSI**D**TNPRAMQA<br>LLQIQQGLQTLQTEAPGLVPSLGSFGISRTPAPSAGSNAGSTPEAPTSSPATPATSSPTGASS<br>AQQQL**K**QQMIQLLAGSGNSQVQTPEVRFQQQLEQLNS**C**G**P**INREANLQAL**P**ATGGDIN<br>A**D**IERLLGSQLS (LEHHHHHH) |
